# Degradation rather than disassembly of necrotic debris is essential to enhance recovery after acute liver injury

**DOI:** 10.1101/2025.01.20.633891

**Authors:** Sara Schuermans, Jusal Quanico, Caine Kestens, Sofie Vandendriessche, Emily Slowikowski, Maria-Laura Crijns, Noëmie Pörtner, Nele Berghmans, Geert Baggerman, Matheus Silvério Mattos, Paul Proost, Pedro Elias Marques

## Abstract

Necrotic cell death causes loss of membrane integrity, release of intracellular contents and deposition of necrotic cell debris. Effective clearance of this debris is crucial for resolving inflammation and promoting tissue recovery. While leukocyte phagocytosis plays a major role, soluble factors in the bloodstream also contribute to debris removal. Our study examined whether enzymatic degradation or disassembly of necrotic debris enhances clearance and improves outcomes in a mouse model of drug-induced liver injury. Using intravital microscopy and proteomic profiling, we demonstrated that necrotic debris is more complex than anticipated, containing DNA, filamentous actin, histones, complement C3, fibrin(ogen) and plasmin(ogen), among many other components. DNase 1 treatment facilitated recovery significantly by enhancing the clearance of DNA from necrotic areas, reducing circulating nucleosomes and actin, and lowering the associated inflammatory response. However, its effect on actin and other damage-associated molecular patterns in necrotic regions was limited. Treatment with short synthetic peptides, specifically 20-amino acid-long positively charged PLK and negatively charged PLE, which displace histones from debris *in vitro,* did not inhibit liver injury or promote recovery. Moreover, activating plasmin to disrupt fibrin encapsulation via tissue plasminogen activator (tPa) led to increased circulating actin levels and worsening of injury parameters. These findings suggest that fibrin encapsulation is important for containing necrotic debris and that enzymatic degradation of necrotic debris is a more effective strategy to enhance tissue recovery than targeting debris disassembly.

**GRAPHICAL ABSTRACT:** 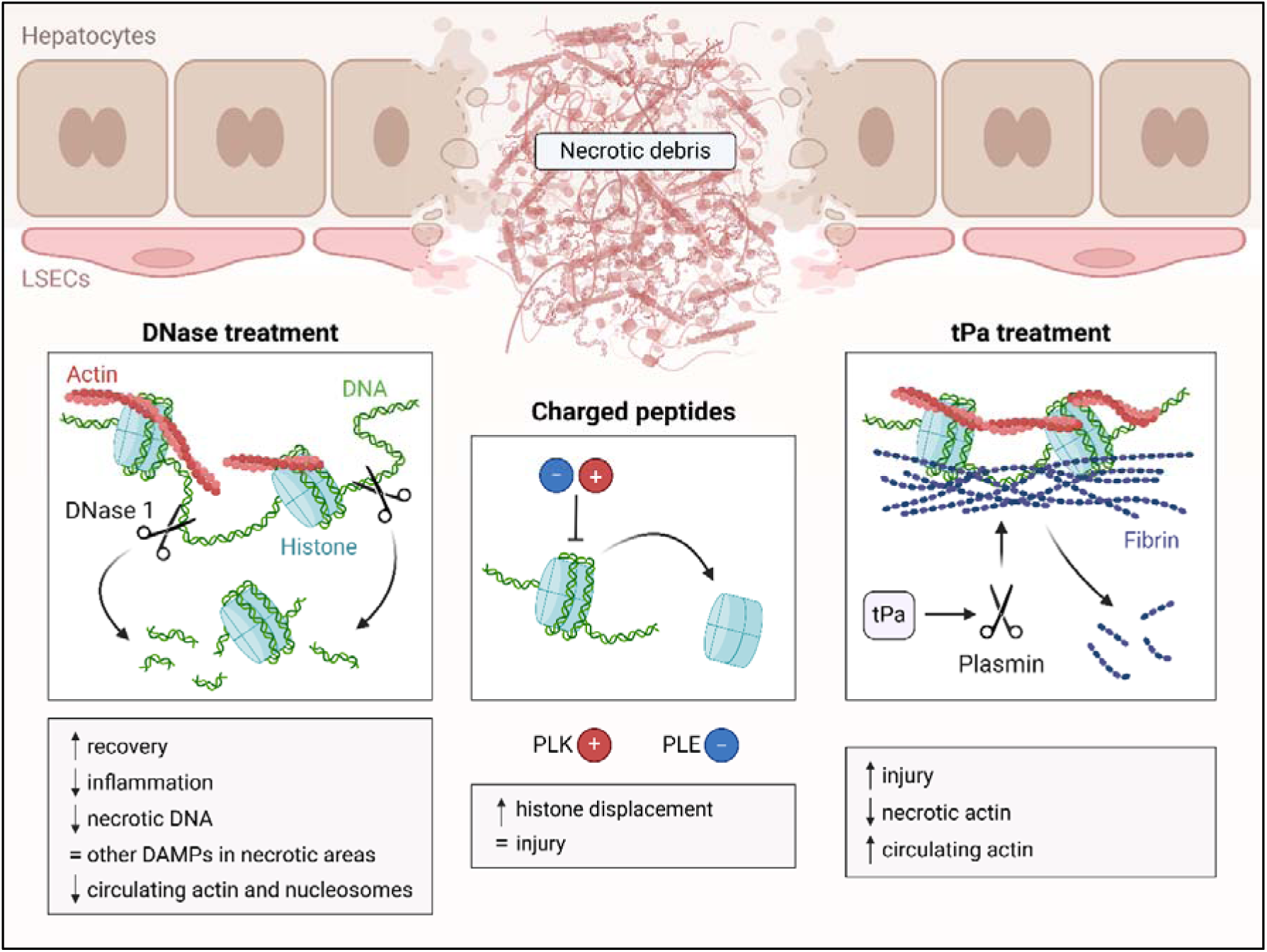

## INTRODUCTION

Cell death is a vital process for maintaining homeostasis, yet it can also be detrimental. Unlike apoptosis, necrosis is characterized by cellular swelling, the irreversible loss of membrane integrity, and the subsequent release of cellular contents, resulting in the formation of necrotic cell debris. This process can occur either accidentally or in a regulated, programmed manner, and is often triggered by extreme external factors such as injury [1]. Necrotic debris primarily consists of damage-associated molecular patterns (DAMPs), which encompass a variety of molecules, including DNA, histones, HMGB1, actin and mitochondria-related molecules. Once exposed, DAMPs trigger inflammation by activating pattern recognition receptors, such as toll-like receptors (TLRs) and C-type lectin receptors (CLECs) [2]. For instance, CLEC2D recognizes nucleosomes by binding to histone tails, carrying the histone-bound DNA complexes into endosomes and stimulating DNA recognition by TLR9 [3]. Extracellular histones also interact with TLR2 and TLR4, an effect intensified when bound to DNA [4]. Additionally, polymerized F-actin released from necrotic cells is detected by CLEC9A (DNGR-1) on dendritic cells [5, 6]. While these inflammatory responses are crucial for tissue repair, unresolved necrotic debris can lead to excessive inflammation and collateral damage. Conditions such as drug-induced liver injury [7, 8], trauma [9] and burn injury [10] highlight the harmful effects of persistent necrotic debris, emphasizing the need for its efficient clearance.

While apoptotic cell clearance (efferocytosis) has been extensively studied, the mechanisms underlying necrotic debris clearance remain less understood. Nevertheless, it is known that necrotic debris clearance involves phagocytosis, which relies on both natural antibodies and complement activation [11, 12]. Beyond phagocytosis by leukocytes, soluble enzymes in the bloodstream play a crucial role in necrotic debris clearance [2]. Among these, deoxyribonuclease (DNase) 1 and DNase 1L3 are essential for degrading necrotic DNA, one of the most potent inflammatory DAMPs [8, 13, 14]. DNase 1 preferentially cleaves double-stranded “naked” linker DNA, while DNase 1L3 efficiently targets complexed DNA, such as chromatin [15–17]. Despite their presence in the circulation, necrotic DNA is often accumulated in injury sites, suggesting that mechanisms take place during necrotic cell death that delay DNA clearance, exacerbating inflammatory responses [12].

Impaired necrotic DNA clearance may stem from electrostatic interactions among necrotic debris components, which increase resistance to degradation. DNA and actin, both negatively charged, can interact with positively charged histones, potentially forming stable complexes that resist degradation [2]. Furthermore, actin inhibits DNase 1 directly by forming complexes with the enzyme, preventing it from degrading necrotic DNA [18, 19]. These challenges highlight the potential benefits of strategies aimed at enhancing DNase activity or disassembling necrotic debris to facilitate its clearance and mitigate inflammation.

Besides the presence of typical DAMPs in necrotic debris, fibrin deposition at injury sites acts as a scaffold for thrombus formation and coagulation. The fibrinolytic system, driven by plasmin activation, balances out the vascular occlusion and promotes injury resolution [20]. However, fibrin deposition may become more resistant to fibrinolysis by interacting with necrotic debris, particularly actin and histones [21–23]. Plasmin has a direct impact on necrotic debris clearance, collaborating with DNase 1 to degrade chromatin by targeting linker histone H1, thereby enhancing DNase 1 access to DNA [24]. Additionally, plasmin-mediated proteolytic cleavage is crucial for degrading necrotic debris both *in vitro* and *in vivo* [25, 26]. Thus, enhancing plasmin activation could improve necrotic debris degradation.

In this study, we used confocal intravital microscopy to observe necrotic debris in the livers of mice following drug-induced liver injury. Our objective was to determine whether enzymatic degradation or disassembly of necrotic debris could enhance its clearance and promote recovery. We hypothesized that DNase 1 treatment would accelerate necrotic DNA degradation, thereby reducing its scaffolding effect and facilitating recovery. Our findings showed that necrotic debris consists largely of extracellular DNA and F-actin. Contrary to our hypothesis, DNase 1 treatment did not affect F-actin levels within the necrotic regions, but it did facilitate recovery, potentially by aiding in the clearance of circulating nucleosomes and actin, and reducing the bulk of DNA debris locally. Proteomic analysis indicated that this treatment caused only minor changes in the protein composition of the necrotic areas and did not significantly alter key DAMPs. Additionally, we investigated whether disassembling necrotic debris would expose it to the nuclease-rich bloodstream, enhancing its degradation. Both positively charged PLK and negatively charged PLE peptides were able to disassemble necrotic debris. While both peptides were effective in displacing histones from necrotic debris *in vitro*, disassembling the debris alone did not improve outcomes *in vivo*. Furthermore, activating plasmin through tissue plasminogen activator (tPa) to disrupt fibrin encapsulation of necrotic debris unexpectedly worsened liver injury. In conclusion, enzymatically degrading necrotic debris, rather than merely disassembling it, is crucial for recovery after drug-induced liver injury.

## RESULTS

### F-actin is found in necrotic areas alongside extracellular DNA and remains intact after DNase injection

To investigate the composition and biochemical properties of necrotic debris *in vivo*, a mouse model of acetaminophen (APAP)-induced liver injury was utilized, which is characterized by extensive hepatocyte necrosis [7, 8]. Mice were administered a sublethal dose of 600 mg/kg APAP via oral gavage, resulting in centrilobular necrosis 24 hours post APAP overdose and recovery by 48 hours [7, 11]. Intravital microscopy of mouse livers revealed necrotic areas marked by extracellular DNA deposition, visualized with SYTOX Green. Fluorescently labeled phalloidin showed the presence of extracellular F-actin in these necrotic areas, alongside the DNA. Interestingly, despite the extensive cell death, the actin cytoskeleton within the dead hepatocytes remained intact (**Figure 1A**). These findings were corroborated using an *in vitro* model of necrotic hepatocyte debris, in which DNA, (F-)actin and histones were all present as well (**Supplementary Figure 1A**). In contrast, attempts to visualize centrilobular necrosis by staining liver cryosections proved inadequate. Staining for DNA (SYTOX) and F-actin (Phalloidin) showed minimal staining in necrotic areas (**Supplementary Figure 1B**). Additionally, immunostaining for G-actin and histone H3 was also absent in these necrotic regions (**Supplementary Figure 1C**). These observations are evidence of the inadequacy of processed tissues sections to investigate necrotic cell debris, justifying the use of confocal intravital microscopy to obtain direct insight into necrotic debris *in vivo*. To examine the stability of necrotic debris *in vivo* and assess whether DNA serves as a scaffold for other debris components, mice were injected with DNase 1 [8]. First, to visualize DNase 1 localization, inactive, fluorescently labeled DNase 1 was injected. It localized specifically to necrotic areas, in contrast to fluorescently labeled bovine serum albumin (BSA) used as a control, which was equally present inside the liver vasculature and the necrotic areas (**Figure 1B, Supplementary Movie 1**). Furthermore, unlabeled, active DNase 1 rapidly degraded approximately 35% of the extracellular DNA in the necrotic areas, however, without affecting F-actin levels at these sites (**Figure 1C, Supplementary Movie 1**). The SYTOX Green signal remained stable over time in the absence of DNase 1 treatment, confirming that the observed reduction in DNA signal was specifically due to DNase 1 activity and not a result of photobleaching (**Supplementary Figure 1D**). These results show that while F-actin is released alongside DNA during necrotic injury, the degradation of DNA by DNase 1 does not alter F-actin locally. Consequently, DNA does not function as a scaffold for actin in necrotic debris. These observations also underscore the importance of using intravital microscopy over immunostaining to accurately investigate necrotic cell debris.

**Figure 1.**
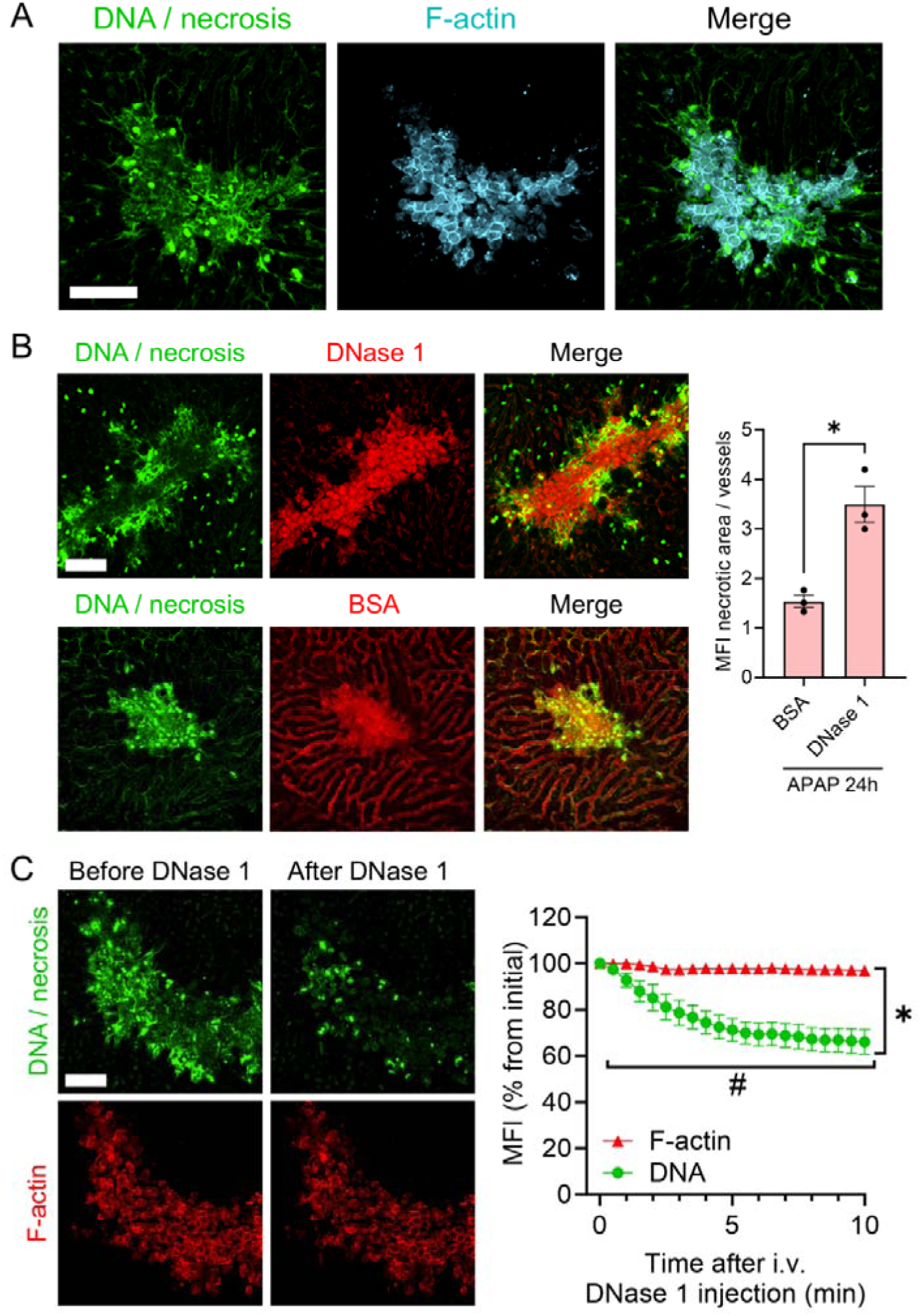
F-actin is found in necrotic areas alongside extracellular DNA and remains intact after DNase injection. **(A)** Representative intravital microscopy images of mice showing deposition of F-actin (Alexa Fluor 555 Phalloidin^+^, 2.2 µM) alongside extracellular DNA (SYTOX Green^+^, 10 µM) in the liver 24 hours after APAP overdose (600 mg/kg). Green: DNA; Cyan: F-actin. **(B)** Representative intravital microscopy images of fluorescently labeled DNase 1 (Alexa Fluor 594, 50 µg) localizing to necrotic areas 24 hours after APAP overdose, compared to fluorescently labeled bovine serum albumin (BSA) as a control. The mean fluorescent intensities (MFIs) of these dyes in the necrotic areas were normalized to the MFIs in the vessels. Data are represented as mean ± SEM. Each dot represents a single mouse. *p≤0.05 between indicated groups. Green: DNA; Red: DNase 1 or BSA. **(C)** Representative intravital microscopy images of mice 24 hours after APAP overdose, showing DNA and F-actin labeling in the liver before and after intravenous (i.v.) DNase 1 injection (40 mg/kg). #p ≤ 0.05 compared to time point 0 for DNA and F-actin. *p ≤ 0.05 between DNA and F-actin at 10 minutes after DNase injection. Data are represented as mean ± SEM. Green: DNA; Red: F-actin. Quantifications were pooled from 3 fields of view per mouse. APAP, acetaminophen. Scale bars = 50 µm.

### Degradation of extracellular DNA by DNase 1 enhances recovery in drug-induced liver injury

Previous studies have demonstrated that systemic DNase 1 treatment reduces liver injury and inflammation 24 hours after an APAP challenge [8], however, the impact of DNase 1 treatment on the recovery phase and the systemic levels of DAMPs was not investigated. Mice challenged with APAP received intravenous (i.v.) injections of DNase 1 (40 mg/kg) 6, 12 and 24 hours post-overdose for evaluation at the 48-hour time point, which was chosen to monitor specifically the recovery phase and the extent of tissue repair [11].

Both serum ALT levels and fibrin(ogen) deposition in the liver were significantly reduced at 48 hours in the DNase-treated mice (**Figure 2A, B, D**). This suggests that the degradation of necrotic DNA enhances recovery after drug-induced liver injury. Conversely, no differences were observed in the expression of Ki67 in liver cryosections, indicating that cellular proliferation was not affected by DNase 1 treatment (**Figure 2C**). Fibrin(ogen) deposition was localized to regions where actin cytoskeleton labeling was lost in tissue sections during necrotic liver injury. As liver regeneration occurs, the actin labeling returns, providing evidence of repair, and fibrin(ogen) deposition diminishes (**Figure 2D**). Moreover, 48 hours after APAP overdose, there was a significantly lower recruitment of neutrophils (40% reduction in the DNase 1-treated group) and a higher recruitment of classical monocytes to livers of DNase 1-treated mice (**Figure 2E, F**). The percentage of recruited non-classical monocytes remained unchanged, and there was a trend toward increased macrophage repopulation in the liver (**Figure 2G, H**). This indicates that, in general, the livers of DNase-treated mice are less inflamed and are further advanced in the process of recovery.

**Figure 2.**
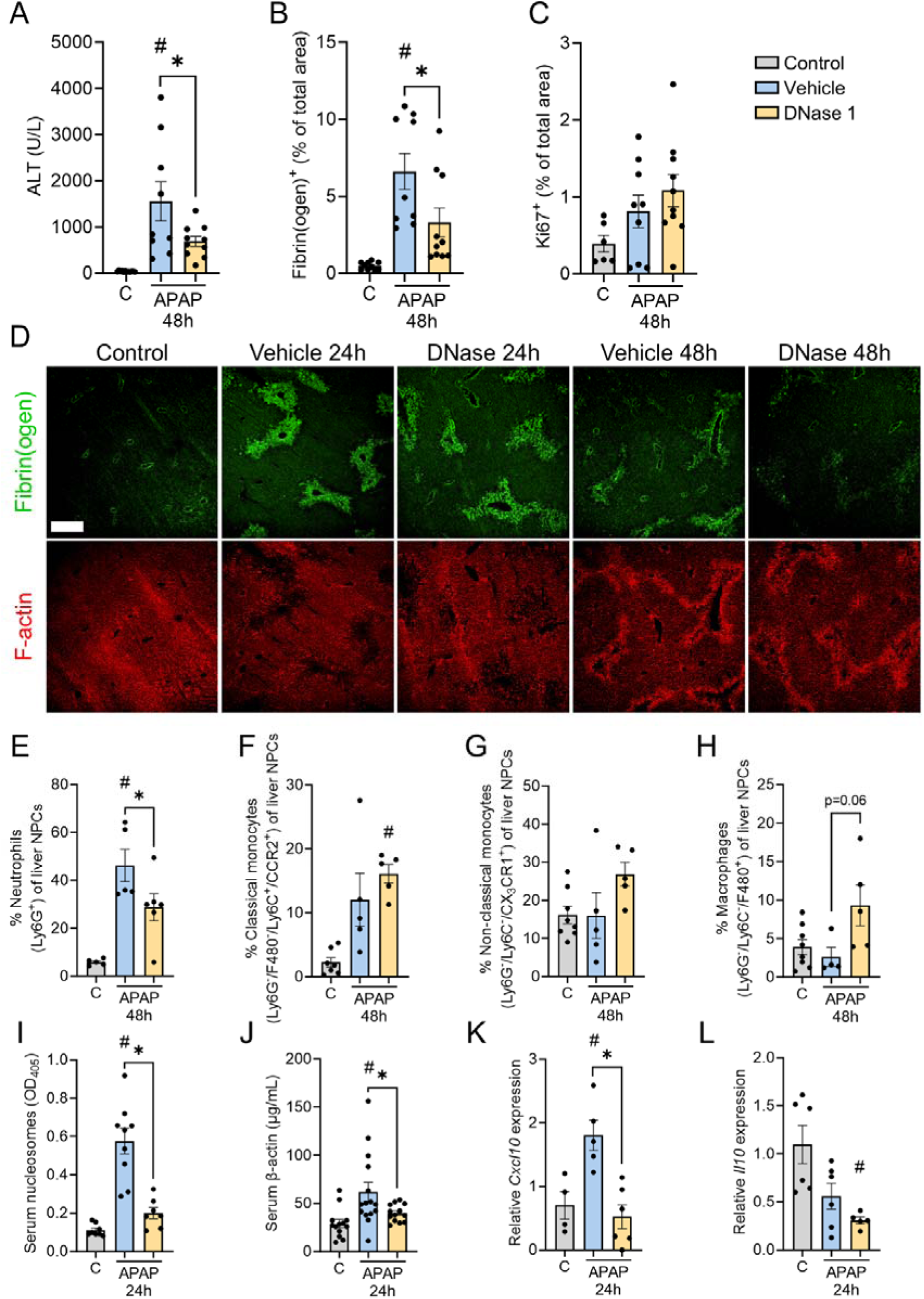
Degradation of extracellular DNA by DNase 1 enhances recovery in drug-induced liver injury. After receiving an APAP overdose (600 mg/kg), mice were sacrificed 24 or 48 hours later for analysis. For the 24-hour time point, mice were treated i.v. with either vehicle (PBS) or DNase 1 (40 mg/kg) 6 and 12 hours post-overdose. For the 48-hour time point, an additional dose of DNase 1 was administered 24 hours after the overdose. For those mice, the following parameters were evaluated at 48 hours: **(A)** Serum alanine aminotransferase (ALT) levels, **(B)** fibrin(ogen)^+^ and **(C)** Ki67^+^ area fraction in liver cryosections. **(D)** Representative images showing immunostaining of liver cryosections from APAP-challenged mice treated with vehicle or DNase 1. Green: fibrin(ogen); Red: F-actin. Scale bar = 200 µm. For liver cryosections, quantifications were pooled from 10 fields of view per mouse. **(E)** Flow cytometry of liver non-parenchymal cells (NPCs) 48 hours post-APAP showing the percentage of neutrophils (Ly6G^+^), **(F)** classical monocytes (Ly6G^+^/F480^−^/Ly6C^+^/CCR2^+^), **(G)** non-classical monocytes (Ly6G^+^/Ly6C^−^/CX3CR1^+^) and **(H)** macrophages (Ly6G^−^/Ly6C^−^/F480^+^). At 24 hours post-APAP, following parameters were evaluated: **(I)** Serum nucleosome and **(J)** serum β-actin levels, **(K, L)** *Cxcl10* and *Il10* expression levels in mouse livers, normalized to the average expression of a housekeeping gene (*Cdkn1a*), and presented as 2^−ΔΔCt^ relative to the control group. Each dot represents a single mouse. Data are represented as mean ± SEM. *p ≤ 0.05 between indicated groups. # p ≤ 0.05 compared to control. C, control; APAP, acetaminophen.

During the peak of injury (24 hours after APAP overdose), an increase in circulating nucleosomes and actin was observed (**Figure 2I, J**). Notably, DNase 1 treatment led to a reduction in both circulating nucleosomes and actin, indicating that DNase 1, through the enzymatic degradation of DNA, enhances the clearance of these components. To further illustrate how these circulating DAMPs are cleared under normal conditions and how they might trigger inflammatory responses, healthy mice were injected i.v. with fluorescently labeled hepatocyte debris. The clearance process by the reticuloendothelial system, including both Kupffer cells and liver sinusoidal endothelial cells (LSECs), was visualized using liver intravital microscopy. Debris in the systemic circulation was primarily taken up by Kupffer cells, although LSECs also contributed to debris uptake. Interestingly, when Kupffer cells were depleted using clodronate liposomes, LSECs did not compensate for this by increasing their debris uptake (**Supplementary Figure 2A, B**). The reduction in circulating DAMPs by DNase treatment was corroborated by decreased expression of the proinflammatory gene *Cxcl10* and the anti-inflammatory gene *Il10* in the livers of DNase-treated mice (**Figure 2K, L**). The levels of *Cxcl1* expression in the liver were not altered across the treatment groups (**Supplementary Figure 2C**). To conclude, DNase 1 treatment improves recovery, potentially by facilitating the clearance of circulating nucleosomes and actin.

### Proteomic analysis of injured liver areas shows the broad composition of necrotic cell debris

To further investigate the composition of necrotic cell debris and broaden the scope of our analysis, and to determine whether DNase 1 treatment alters the debris *in vivo*, bottom-up proteomics was performed on the necrotic areas of mice that received an APAP overdose, followed by treatment with either vehicle or DNase 1. The analysis was conducted at the 24-hour timepoint, coinciding with the peak of injury when substantial debris had accumulated in the necrotic regions. **Figure 3A** depicts a heatmap with two distinct clusters of proteins: the upper cluster consists of proteins enriched in the necrotic areas of vehicle-treated mice, while the lower cluster contains proteins enriched in the DNase-treated group. Proteins detected and quantified in at least 30% of events within a given experimental group were retained, resulting in 936 proteins that met the criteria and were selected for further analysis. Of these, only 42 proteins (4%) showed significant alterations between groups. Notably, DNase 1 treatment did not affect the composition of key DAMPs such as histones, actin and HMGB1, suggesting that a component other than DNA may serve as a scaffold for the necrotic debris. Other proteins detected but not significantly altered between the two conditions included components of the coagulation, fibrinolytic and complement systems, such as fibrinogen, complement C3 and plasmin(ogen). **Figure 3B** presents a volcano plot highlighting the proteins that were significantly altered, using a twofold threshold for increases or decreases. On the right side of the plot are proteins that were enriched in the necrotic areas of DNase-treated mice, while the left side displays proteins more abundant in the necrotic areas of vehicle-treated mice. Proteins that were enriched in the DNase-treated mice and were annotated in the volcano plot are: inactive rhomboid protein 1 (RHDF1), arylacetamide deacetylase (AAAD), N-acyl-aromatic-L-amino acid amidohydrolase (ACY3), large ribosomal subunit protein uL10 (RLA0), zyxin (ZYX) and small ribosomal subunit protein RACK1 (RACK1). The proteins that were enriched but were not annotated in the figure include: sulfotransferase 1A1 (ST1A1), spermatogenesis-associated protein 20 (SPT20), TFG, cytochrome P450 2J5 (CP2J5), interleukin enhancer-binding factor 2 (ILF2) and long-chain-fatty-acid--CoA ligase 5 (ACSL5). The proteins that were enriched in the vehicle-treated group include: cytochrome P450 2D9 (CP2D9), BAG family molecular chaperone regulator 3 (BAG3), microsomal glutathione S-transferase 1 (MGST1), rho GDP-dissociation inhibitor 1 (GDIR1), apolipoprotein A-II (APOA2), 55 kDa erythrocyte membrane protein (EM55) and cytoplasmic dynein 1 light intermediate chain 1 (DC1L1). Of note, the latter proteins are potential biomarkers and could be used as indicators of liver necrosis in biopsies in the clinic. No gene ontology enrichment analysis was performed, as the proteins originated from necrotic debris, and thus dead cells, making such analysis irrelevant. Furthermore, no clear link among these various proteins has been identified. In conclusion, our data demonstrates that necrotic cell debris is vastly complex and composed by both cellular and extracellular molecules. Moreover, DNase 1 treatment induces minimal changes in protein composition within necrotic regions and does not affect the levels of key DAMPs such as histones, actin or HMGB1.

**Figure 3.**
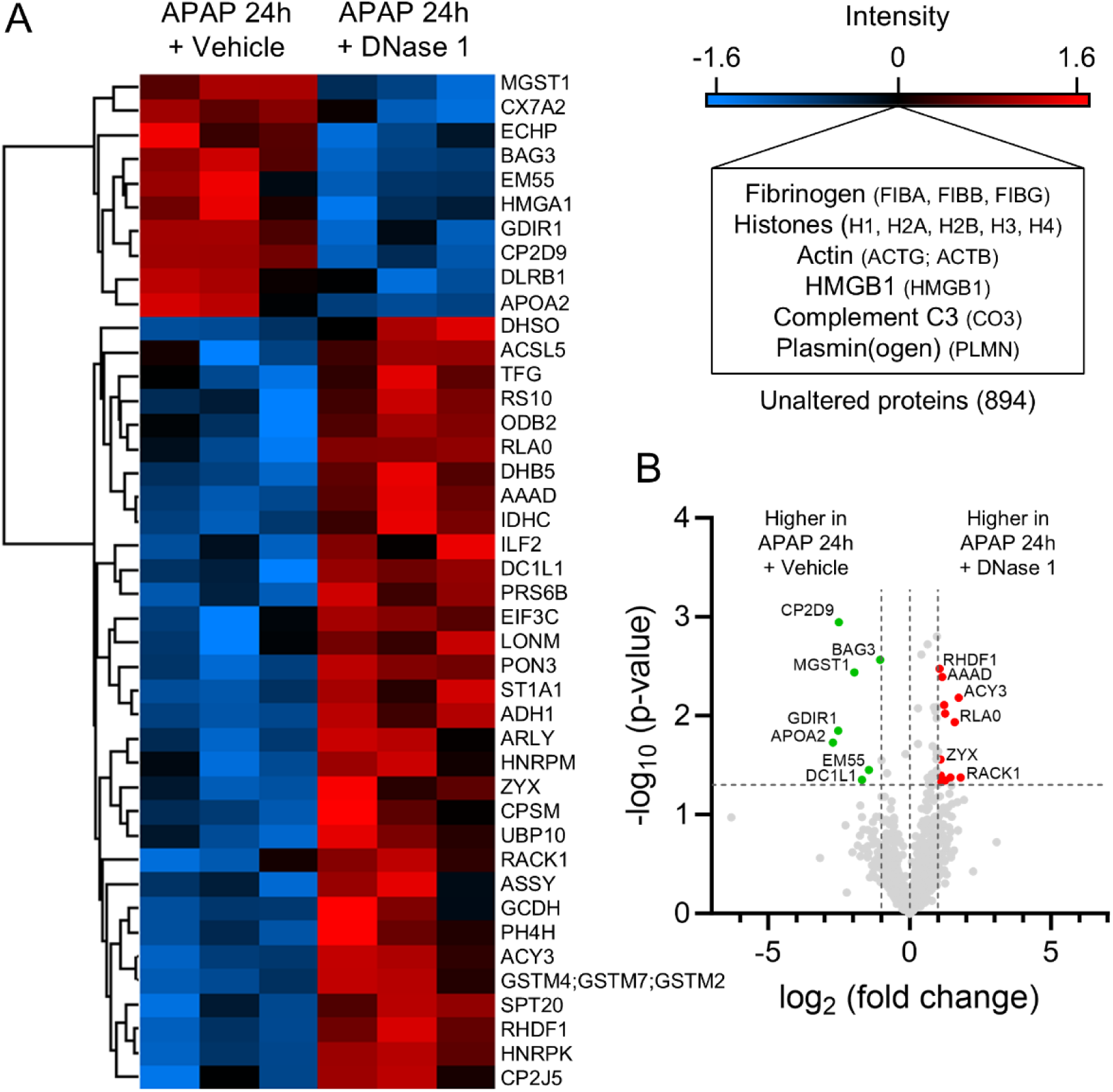
Proteomic analysis of necrotic areas after DNase 1 treatment. Mice received an APAP overdose (600 mg/kg), were treated i.v. with either vehicle (PBS, n = 3) or DNase 1 (40 mg/kg, n = 3) at 6 and 12 hours post-overdose, and were sacrificed at 24 hours. Cryosections were made, and on-tissue microdigestion with trypsin of injured areas was followed by identification using LC-MS/MS. **(A)** Heatmap of differentially expressed proteins in necrotic areas between mice that were treated with vehicle or DNase 1. LFQ intensities were log2-transformed and median-normalized row-wise, with higher z-scores and thus higher relative abundances for each protein indicated in red. Each row represents a protein, each column is a sample. Comparisons with p≤0.05 were considered significant. Proteins that were detected but not significantly altered between both conditions included DAMPs and proteins of the coagulation, fibrinolytic and complement system. **(B)** Volcano plot of the differentially expressed proteins, showing the fold change (log_2_ ratio) plotted against the statistical significance (-log_10_ of the p-value). Red and green dots in volcano plot indicate proteins with significant differences (green, p ≤ 0.05, 2-fold decrease; red, p ≤ 0.05, 2-fold increase); gray dots are proteins without significant change. APAP, acetaminophen.

### Charged peptides displace debris *in vitro* but do not improve outcome *in vivo*

Interactions within necrotic debris might be charge-dependent, potentially involving DNA, histones and actin. The disassembly of DNA-protein complexes could allow serum DNases to access and digest the extracellular DNA more effectively, therefore, we investigated whether these interactions could be disrupted using positively and negatively charged peptides. The peptides used in this study included PLK and PDK, both positively charged and composed of 20 lysine residues with L- or D-amino acids respectively, and PLE, a negatively charged peptide composed of 20 glutamate residues. These peptides were chemically synthesized and purified (**Supplementary Figure 3A, B**). Necrotic hepatocyte debris was generated *in vitro* by mechanical disruption of HepG2 cells, and the resulting debris was then incubated with the peptides of interest. PLK successfully displaced histones from necrotic debris at concentrations ranging from 1 µM to 10 µM. However, at a concentration of 100 µM, PLK caused a reduction in the amount of actin displaced compared to buffer alone, suggesting that PLK precipitated actin at its highest concentration (**Figure 4A**). The D-amino acid variant, PDK, was ineffective at displacing histones or actin from necrotic debris, but similarly to PLK, it caused actin precipitation at the highest concentration (**Supplementary Figure 4A**). Poly-L-Lysine, a longer polymer used as control for PLK, was also unable to displace histones or actin *in vitro* (**Supplementary Figure 4B**). In contrast, PLE effectively displaced histones from necrotic debris at 100 µM, leading to a tenfold increase in histone displacement compared to buffer alone (**Figure 4B**). Heparin, a highly negatively charged polysaccharide used as a control for PLE, showed a similar effect in displacing histones at its highest concentration, though less pronounced. Heparin also displaced actin at concentrations ranging from 250 U/mL to 500 U/mL (**Supplementary Figure 4C**). To conclude, both positively charged PLK and negatively charged PLE are able to displace histones from necrotic debris *in vitro*.

**Figure 4.**
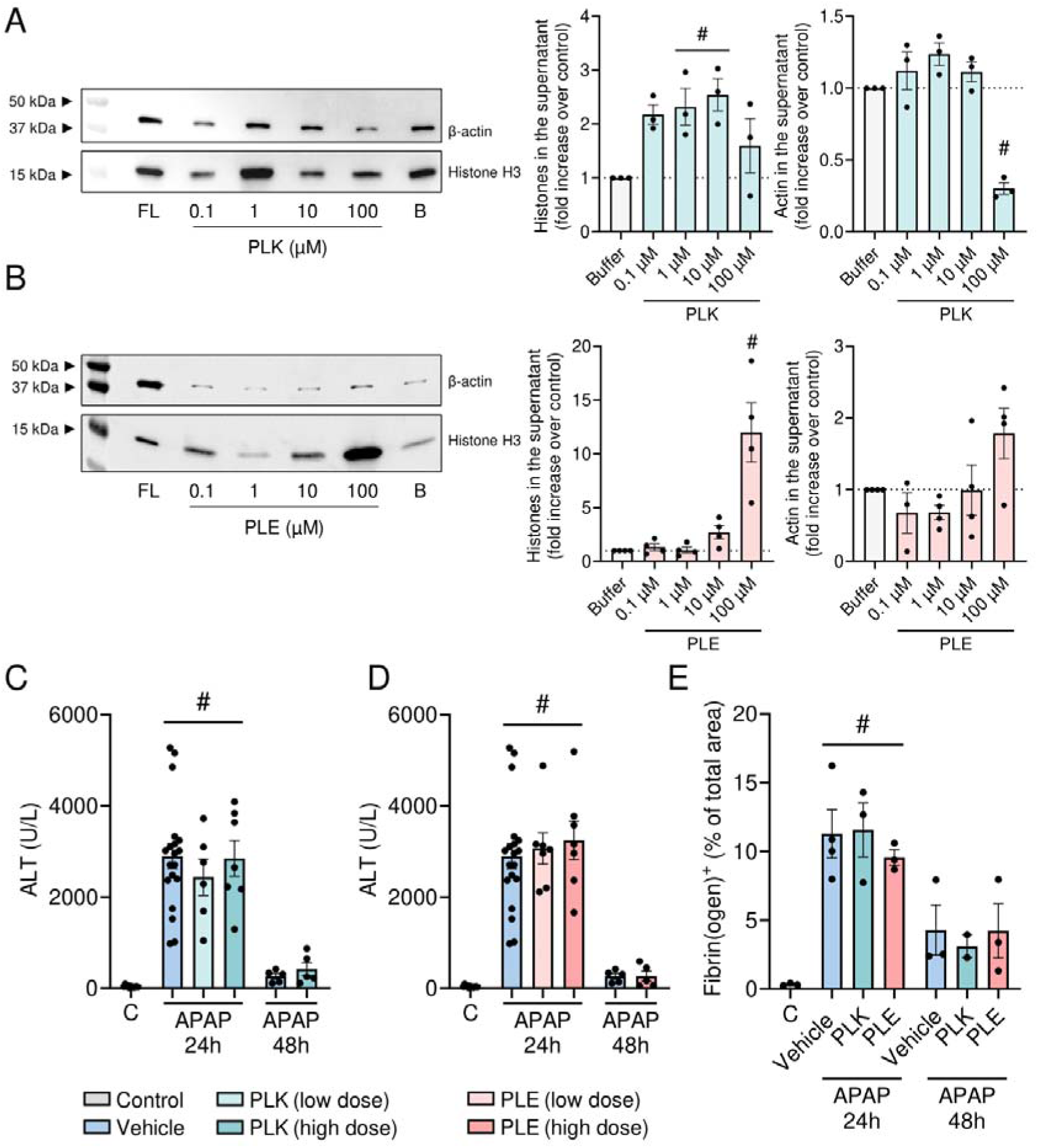
Charged peptides displace debris *in vitro* but do not improve outcome *in vivo*. Western blotting for histone H3 and β-actin in supernatant of HepG2 debris incubated with different concentrations of **(A)** positively charged PLK and **(B)** negatively charged PLE, or buffer (B) alone. A full lysate of HepG2 cells (FL) was loaded at 10 µg as a positive control. Mice that received an APAP overdose (600 mg/kg) were treated i.v. with 400 µg/kg (low dose) or 800 µg/kg (high dose) of the peptides at 6 and 12 hours for the 24-hour time point, with an additional dose given at 24 hours for the 48-hour time point. **(C, D)** Serum alanine aminotransferase (ALT) levels following treatment with PLK or PLE. **(E)** Fibrin(ogen)^+^ area fraction in liver cryosections. Quantifications for liver cryosections were pooled from 10 fields of view per mouse. Each dot represents a single mouse. Data are represented as mean ± SEM. #p ≤ 0.05 compared to control. C, control; APAP, acetaminophen.

Next, the peptides were tested *in vivo* to assess whether they could disrupt necrotic debris and improve outcomes in mice suffering from drug-induced liver injury. Mice treated twice with PLK, PDK or PLE at both low and high doses (400 µg/kg and 800 µg/kg) showed no differences in serum ALT levels at 24 hours, nor after being treated three times and analyzed at 48 hours post-overdose, indicating no protective effect from the peptides (**Figure 4C, D; Supplementary Figure 4D**). Additionally, immunostaining of liver cryosections for fibrin(ogen), F-actin and histone H3 was performed (**Supplementary Figure 5**). The fibrin(ogen) area fraction in the liver was unchanged at both 24 and 48 hours after the APAP overdose, even with the highest doses of PLK and PLE (**Figure 4E**). This shows that histone-displacing peptides do not improve outcomes in mice with drug-induced liver injury.

### Increasing fibrinolysis worsens outcome after drug-induced liver injury

Fibrin deposition is a hallmark of injury and was proposed to hinder the degradation of necrotic cell debris [2]. Considering that debris-dissociating peptides were insufficient to ameliorate liver injury, we hypothesized that fibrin may be the key factor anchoring the debris in place. To investigate whether the accumulation of debris in the liver contained by the fibrin capsule might be detrimental, fibrinolysis was stimulated using tissue plasminogen activator (tPa). Intravital microscopy of livers 24 hours post-APAP overdose showed that treatment with tPa removed approximately 10% of F-actin from necrotic areas *in vivo* within 30 minutes, without altering extracellular DNA levels (**Figure 5A, Supplementary Movie 2**). Mice were also challenged with APAP for 24h and received i.v. injections of tPa (5 mg/kg) at 12 hours post-overdose. In tPa-treated mice, a larger area of the liver was covered in fibrin(ogen), corresponding to an expanded region of necrotic liver injury, as quantified in cryosections (**Figure 5B, D**). No differences in serum ALT levels were observed (**Figure 5C**), however, the worsening of liver injury was corroborated by an increased release of actin into the circulation in tPa-treated mice (**Figure 5E**). The expression levels of the proinflammatory genes *Cxcl1* and *Cxcl10* in the livers of tPa-treated mice were reduced compared to vehicle-treated mice, while no differences in *Il10* expression were observed (**Figure 5F, G; Supplementary Figure 2D**). Overall, treatment with tPa to stimulate fibrinolysis appeared to worsen the condition of APAP-treated mice, likely by increasing the release of debris into the circulation. Additionally, larger necrotic areas were observed, possibly due to the early disruption of the fibrin barrier that covered the injured vasculature and tissue, resulting in increased injury. To further confirm that the disruption of the interaction between fibrin and the necrotic debris, leading to debris release into the bloodstream, contributes to worsening of liver injury, mice were treated 6 hours after APAP overdose with either heparin [3000 U/kg, intraperitoneally (i.p.)] or actin (800 µg/kg i.v.). Inhibition of coagulation by heparin resulted in higher serum ALT levels 24 hours post-APAP overdose, indicating that fibrin deposition on necrotic debris may contain liver injury. Similarly, actin injection to emulate more debris being released significantly elevated serum ALT levels, suggesting that the release of DAMPs into circulation is directly harmful (**Figure 4H**). Together, these findings indicate that interfering with the fibrin encapsulation of necrotic debris exacerbates drug-induced liver injury in mice.

**Figure 5.**
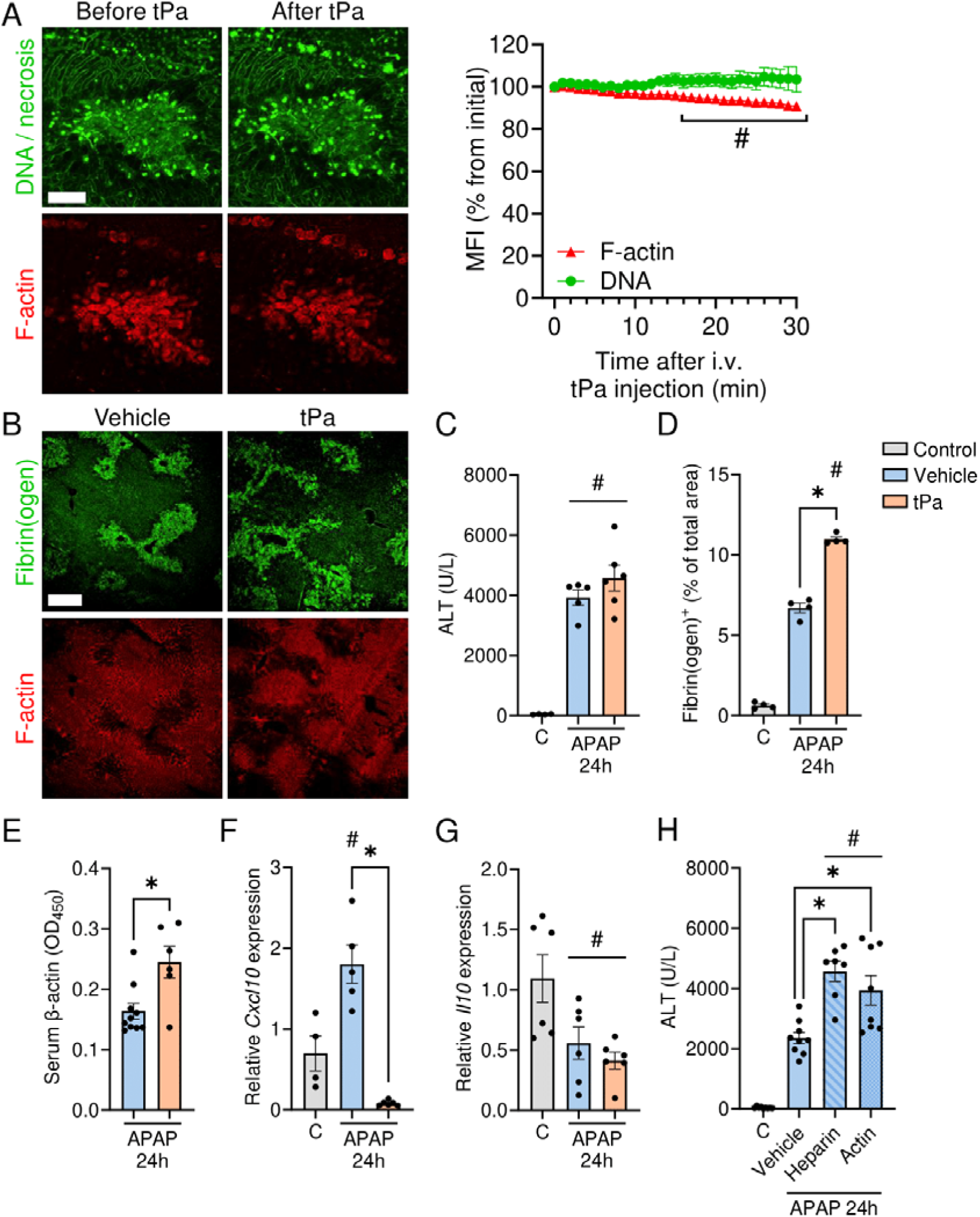
Increasing fibrinolysis worsens outcome after drug-induced liver injury. **(A)** Representative intravital microscopy images of mice 24 hours after APAP overdose (600 mg/kg), showing DNA and F-actin labeling in the liver before and after i.v. tPa injection (5 mg/kg). #p ≤ 0.05 compared to time point 0 for DNA and F-actin. Green: DNA; Red: F-actin. Quantifications were pooled from 3 fields of view per mouse. Scale bar = 50 µm. **(B)** Mice were treated with vehicle (PBS) or tPa (5 mg/kg) 12 hours after receiving an APAP overdose, then sacrificed 24 hours post-APAP. Representative images showing immunostaining of liver cryosections from APAP-challenged mice treated with vehicle or tPa. Green: fibrin(ogen); Red: F-actin. Scale bar = 200 µm. Additionally, the following parameters were assessed: **(C)** Serum alanine aminotransferase (ALT) levels, **(D)** fibrin(ogen)^+^ area fraction in liver cryosections, with quantifications pooled from 10 fields of view per mouse, **(E)** serum β-actin levels and **(F-G)** *Cxcl10* and *Il10* expression levels in mouse livers, normalized to the average expression of a housekeeping gene (*Cdkn1a*), and presented as 2^−ΔΔCt^ relative to the control group. **(H)** Serum ALT levels in mice treated with heparin i.p. (3000 U/kg) or actin i.v. (800 µg/kg) 6 hours after APAP overdose and sacrificed at 24 hours. Each dot represents a single mouse. Data are represented as mean ± SEM. *p ≤ 0.05 between indicated groups. # p ≤ 0.05 compared to control. C, control; APAP, acetaminophen.

## DISCUSSION

Although several cellular components and DAMPs have been proposed to be exposed during necrotic cell death, few have been actually demonstrated and quantified *in vivo*. In our research, using a mouse model of drug-induced liver injury, we not only confirmed the presence of extracellular DNA in necrotic regions [7, 8], but also identified the presence of polymerized F-actin. Interestingly, despite membrane permeabilization, the cytoskeletal F-actin within the dead hepatocytes remained intact, indicating that its polymerized form may be important for the recognition of actin as DAMP. Indeed, F-actin associated with specific actin-binding proteins was identified as a ligand for CLEC9A (DNGR-1), rather than its monomeric form [5, 6]. One explanation for why actin remains in its polymerized form could be that F-actin is not digested by proteolytic enzymes, possibly due to the presence of protease inhibitors in plasma. F-actin might also persist in necrotic areas due to the fibrin clot, which would serve both to stabilize and to shield the debris from degradation. Furthermore, fluorescently labeled DNase 1 localized specifically to necrotic regions, indicating its targeted action on necrotic debris. In fact, due to the high affinity between G-actin and DNase 1 [18, 19], fluorescent DNase 1 conjugates are commonly used to detect monomeric G-actin. Therefore, our findings demonstrated that both G- and F-actin are components of necrotic areas *in vivo*.

Regarding the use of intravital microscopy over immunostaining on tissue sections for visualizing necrotic debris *in vivo*, we observed that cryosections were not optimal for imaging debris, however, staining for fibrin(ogen) and the proliferation marker Ki67 was still feasible. This suggests that the issue is specific to detection of debris itself rather than a general limitation of immunostaining in necrotic areas. The debris is partially lost during processing steps or blocked (*e.g.* by fibrin), as DNA, actin and histone H3 were affected, while fibrin(ogen) was not. This argument is substantiated by the observation that all components were present in the hepatocyte debris spot and detected during the proteomic analysis, the latter a label-free technique.

To examine the stability of necrotic debris *in vivo* and determine whether DNA serves as a scaffold for other debris components, mice were injected with DNase 1 and monitored using intravital microscopy [8]. Although DNase 1 injection reduced extracellular DNA in liver necrotic regions, it did not affect F-actin levels. Under healthy conditions, DNase 1 effectively depolymerizes actin filaments through a dual mechanism: inhibiting polymerization due to its high affinity binding to G-actin and promoting F-actin depolymerization by destabilizing the filaments [27]. However, during drug-induced liver injury, the abundant release of actin likely leads to a reduction in DNase 1 activity [9]. Bottom-up proteomics of necrotic areas 24 hours post-APAP overdose revealed no differences in the presence of several DAMPs after multiple DNase 1 treatments, including actin. Since this is a label-free method, stabilization by phalloidin labeling cannot account for the persistence of F-actin in these areas [27], suggesting again that the fibrin clot may be the culprit.

Although DNase 1 had a minor effect on DAMPs other than DNA within necrotic regions, multiple doses of DNase 1 reduced circulating levels of both nucleosomes and actin. DNase 1 digests linker DNA, generating mono- and oligonucleosomes detectable by the nucleosome ELISA kit. Similarly, in a mouse model of myocardial infarction, DNase 1 treatment reduced plasma nucleosome levels 6 hours post-infarction [28]. This reduction may result from DNase 1 fragmenting chromatin into smaller nucleosome units, promoting clearance by the reticuloendothelial system. Indeed, when we injected fluorescently labeled debris into healthy mice, the reticuloendothelial system efficiently took up this debris. The reduction in circulating actin levels may result from DNase 1-mediated depolymerization of F-actin, enhancing its removal. Additionally, degradation of extracellular DNA by DNase 1 could disrupt DNA-actin complexes in circulation, further promoting actin clearance. While DNA-actin complexes have not yet been detected in circulation, they have been identified in the sputum of cystic fibrosis patients where DNase 1 is used therapeutically to reduce viscosity [29]. Overall, DNase 1 treatment enhanced recovery from drug-induced liver injury and mitigated the associated inflammatory response.

To uncover overlooked molecules deposited in necrotic sites, a proteomic analysis was performed to assess the necrotic debris composition in the liver. Repeated DNase 1 treatments following APAP overdose did not affect the majority of debris components at peak injury, reinforcing the notion that multiple simultaneous mechanisms are required to clear debris from necrotic regions. Regarding the differences in specific proteins between conditions, further studies are needed to explore its implications. In general, proteomic studies on necrotic regions have not yet been conducted, leaving no basis for direct comparison. However, our findings indicate the necessity of protease-containing processes and phagocytes for the degradation of necrotic debris. Neutrophils, which are among the first immune cells recruited to necrotic regions [30], contain matrix metalloproteinases (MMP-9) and serine proteases (*e.g.* neutrophil elastase). MMP-9 targets a myriad of DAMPs, including actin, HMGB1 and heat shock protein 90 [31]. Besides this, plasmin is a major proteolytic enzyme that is already implicated in the clearance of debris [24–26].

We also considered the possibility that charge-dependent interactions within necrotic debris might be of importance. It has already been demonstrated that DNA and F-actin bundles are formed upon addition of histone H1 [32]. Therefore, we investigated whether these interactions could be disrupted using positively and negatively charged peptides. The development of peptides for disrupting necrotic debris was based on MIG30, a 30-amino acid long C-terminal fragment of CXCL9 [CXCL9(74-103)], a highly positively charged peptide that competes with chemokines for binding to glycosaminoglycans [33] and protects against liver ischemia-reperfusion injury [34] and drug-induced liver injury in mice [7]. MIG30 has been shown to interact with DNA in a charge-dependent manner and bind to DNA in the necrotic regions *in vivo*. Furthermore, 10 µM MIG30 displaced histones when added to necrotic debris *in vitro*, supporting the notion that it disrupts interactions within the necrotic debris [7]. Both positively charged PLK and negatively charged PLE displaced histones from necrotic debris *in vitro*, similar to MIG30. In contrast, PDK did not exhibit this capability, suggesting that the specific conformation of amino acids may play a critical role. Additionally, heparin, a negatively charged polymer, was able to displace both histones and actin from necrotic debris *in vitro*. This raised the possibility that displacing actin may require a larger molecule, given that actin itself is a long polymer. In the sputum of cystic fibrosis patients, polyanions such as poly-aspartate or poly-glutamate disrupt F-actin and DNA complexes. Furthermore, the addition of poly-aspartate increased DNase activity in samples containing DNA bundles formed with histone H1 [32]. This study and our data suggest that disassembling the bundles within necrotic debris could enhance DNase activity, thereby facilitating debris clearance. However, despite the effectiveness of the peptides in displacing necrotic debris *in vitro*, *in vivo* analysis revealed no significant changes in liver injury or recovery. Disassembling necrotic debris alone proved insufficient to improve outcomes in the drug-induced liver injury model, suggesting that additional factors, such as fibrin deposition, may anchor the debris in place and that this event has physiological value.

To explore whether fibrin surrounding necrotic debris contributes to its accumulation, fibrinolysis was stimulated using tissue plasminogen activator (tPa). A small reduction in the F-actin levels in the liver necrotic regions was observed, while no differences in extracellular DNA levels were detected. Moreover, mice treated with a single dose of tPa exhibited significantly worse outcomes, and importantly, showed an increase in circulating actin levels. This suggested that disruption of fibrin by plasmin leads to the release of actin into the circulation. Actin, when bound to fibrin, can stabilize the fibrin complex and delay fibrinolysis [21, 35], but upon the removal of fibrin, actin is released, which may worsen damage and inflammation. Larger fibrin-positive areas were observed in the mice treated with tPa, which may seem contradictory. However, given that there was a 12-hour interval between tPa treatment and sample collection, this time frame allowed for new coagulation to occur. The disruption of the fibrin barrier surrounding the necrotic debris may have initially increased injury, and the subsequent formation of fresh fibrin deposits in the necrotic regions could explain the observed larger fibrin-positive areas.

APAP-induced liver injury activates both coagulation and fibrinolysis [36], which may contribute to its pathology. However, the role of plasmin activation remains unclear, as both insufficient and excessive fibrinolysis can be harmful. For instance, fibrin(ogen) deficiency delays liver repair following APAP overdose in mice [37], indicating a role for fibrin in tissue regeneration or debris containment. Conversely, plasminogen deficiency reduces susceptibility to APAP-induced liver toxicity, suggesting that excessive fibrinolysis can exacerbate injury [38]. Adding to this complexity, actin released from necrotic cells has been reported to accelerate plasmin generation [39], thereby further promoting fibrinolysis and potentially exacerbating injury. Plasminogen activators contribute to liver injury independently of fibrin(ogen) [36] but also aid in clearing necrotic debris [24–26], highlighting the complex and context-dependent effects of plasmin activation. Our study demonstrates that activating plasmin through tissue plasminogen activator (tPa) to disrupt fibrin encapsulation unexpectedly aggravated liver injury. Similarly, while heparin pretreatment reduced coagulation and liver damage [40], its posttreatment led to worse outcomes, underscoring the stabilizing role of fibrin in maintaining necrotic debris containment and limiting damage. Supporting this, i.v. actin injection exacerbated injury, indicating that the disordered release of DAMPs into circulation is harmful. These findings indicate that disrupting fibrin encapsulation of necrotic debris worsens drug-induced liver injury in mice.

In conclusion, our study provides valuable insights into the complex dynamics of necrotic debris during recovery from drug-induced liver injury. Using intravital microscopy, we confirmed the presence of F-actin in necrotic liver areas along with extracellular DNA. Moreover, we emphasize the importance of degrading DAMPs such as DNA via DNases to facilitate recovery. Our findings also suggest that disrupting the fibrin stabilization of necrotic debris or targeting electrostatic interactions within it is detrimental to recovery. Ultimately, enzymatic degradation of necrotic debris, rather than mere disassembly, is crucial for effective recovery, providing important directions for future therapies that aim to increase necrotic cell debris removal.

## MATERIALS & METHODS

### Cell lines

HepG2 cells were cultured at 37 °C and 5% CO_2_ atmosphere in high glucose Dulbecco’s Modified Eagle Medium (DMEM) with GlutaMAX (Gibco, Thermo Fisher Scientific, Waltham, MA, USA), supplemented with 10% FBS (Sigma-Aldrich, Merck, Darmstadt, DE), 20 mM HEPES buffer solution (Gibco) and 1X MEM Non-Essential Amino Acids Solution (Gibco).

### Mice

Male C57BL/6J mice were purchased from Janvier Labs. All mice used in this study were between 8-12 weeks old. The mice were housed in acrylic filtertop cages (5 mice per cage) with an enriched environment (bedding, toys and small houses) at the Animal Facility of the Rega Institute (KU Leuven, BE). Filtered water and food were provided *ad libitum*, and the mice were maintained in a 12-hour dark-light cycle at 21 °C. All experiments were approved and performed following the guidelines of the Ethical Committee for Animal Experiments from KU Leuven (registry number: P128/2021).

### Acetaminophen (APAP)-induced liver injury model

Mice were fasted for 12 hours before receiving a single oral gavage of either PBS or APAP (600 mg/kg, Sigma-Aldrich) dissolved in warm PBS, ensuring complete APAP absorption and enhancing experimental reproducibility. After 24 or 48 hours, mice were euthanized under anesthesia with ketamine (80 mg/kg) and xylazine (4 mg/kg), after which the liver and blood were collected. For peptide treatments, mice were injected i.v. with 400 µg/kg (low dose) or 800 µg/kg (high dose) of peptide twice, administered at 6 and 12 hours after APAP overdose for the 24-hour time point, and an additional dose given at 24 hours for the 48-hour time point. For DNase treatments, mice were injected i.v. with 40 mg/kg DNase 1 (Roche, Basel, CH) at 6 and 12 hours after APAP overdose for the 24-hour time point, with an additional dose administered at 24 hours for the 48-hour time point. For tPa, mice received a single i.v. dose of 5 mg/kg tPa [Actilyse (alteplase), Boehringer Ingelheim, Brussels, BE] 12 hours after APAP overdose and were sacrificed at the 24-hour time point. Actin (800 µg/kg, rabbit skeletal muscle, Cytoskeleton, Denver, CO, USA) was injected i.v. and heparin (3000 U/kg, LEO Pharma, Amsterdam, NL) was administered i.p. Both were administered 6 hours after APAP overdose to mice sacrificed at the 24-hour time point.

### Peptides synthesis and purification

The peptides PLK, PDK and PLE (composed of 20 consecutive L-lysine, D-lysine and L-glutamate residues, respectively) were chemically synthesized by solid-phase peptide synthesis on a P11 peptide synthesizer (Activotec, Cambridge, UK) using fluorenylmethoxycarbonyl (Fmoc) chemistry, as previously described [41]. Briefly, amino acids were sequentially coupled starting from the COOH-terminus toward the NH_2_-terminus after initial coupling to a rink amide resin (Activotec). Peptide Fmoc deprotection was performed with 20% piperidine in N-methyl-2-pyrrolidone (Biosolve, Valkenswaard, NL) and the next amino acid was activated with 0.45 M 2-(1H-benzotriazol-1yl)-1,1,3,3-tetramethyluroniumhexafluorophosphate (Activotec) and 0.45 M 1-hydroxybenzotriazole (Acros Organics, Thermo Fisher Scientific) in dimethylformamide (Acros Organics). UV absorbance was monitored to ensure adequate deprotection before each coupling step. Peptides were cleaved from resins in a solution of 95% trifluoroacetic acid (TFA, Biosolve) in ultrapure water, and they were filtered and washed in tert-butyl methyl ether (Honeywell Riedel-de-Haën, Seelze, DE). The peptides were then purified using reversed-phase high-performance liquid chromatography (RP-HPLC) with a PepMap C18 column (VDS optilab, Berlin, DE) on a Waters 600 pump and Waters 600 controller (Milford, MA, USA). The peptides were eluted by a gradient of 0 to 80% acetonitrile (ACN) in 0.1% TFA, under continuous monitoring by directing 2% of the eluate to an electrospray ionization ion trap mass spectrometer. Fractions containing peptide with the correct relative molecular mass were collected.

### Alanine aminotransferase (ALT) assay

Liver injury was indirectly assessed by monitoring levels of serum ALT utilizing the Infinity™ ALT (GPT) Liquid Stable Reagent (Thermo Fisher Scientific), which was conducted in accordance to the manufacturer’s instructions. Briefly, blood samples were harvested and centrifuged for 10 minutes at 2400 x g and then serum was harvested. Pure serum and two different dilutions (1:5 and 1:20) were added to a 96-well plate, after which the reagent was added at 37 °C. The rate of decrease in absorbance at 340 nm due to the oxidation of NADH to NAD^+^ was measured every 30 seconds (CLARIOstar Plus, BMG Labtech, Ortenberg, DE).

### Nucleosome and actin ELISA

Serum nucleosome and actin levels were measured using an enzyme-linked immunosorbent assay (ELISA). The nucleosome levels were measured using the Cell Death Detection ELISA^PLUS^ (Roche) that allows for the detection of histone-complexed DNA fragments (mono- and oligonucleosomes). Serum β-actin levels were quantified using the PathScan® Total β-Actin Sandwich ELISA Antibody Pair (Cell Signaling Technology, Danvers, MA, USA) and actin protein [(>99% pure): rabbit skeletal muscle, Cytoskeleton] as a standard. Both assays were performed in accordance to the manufacturer’s instructions.

### Gene expression (qPCR)

The liver (10 mg of the caudate lobe) was homogenized in tubes containing 2.8 mm zirconium oxide beads using the Precellys® 24 Tissue Homogenizer (Bertin Technologies, Montigny-le-Bretonneux, FR). Total RNA was extracted by lysing the cells in the presence of β-mercaptoethanol and using the RNeasy® Plus Mini Kit (Qiagen, Hilden, DE) according to the manufacturer’s instructions. Following extraction, total RNA quality and quantity were assessed using a NanoDrop® ND-1000 spectrophotometer (Isogen Life Science, Utrecht, NL), and 1 µg RNA was used for reverse transcription with the High-Capacity cDNA Reverse Transcriptase Kit (Applied Biosystems, Thermo Fisher Scientific). 50 ng of the resulting cDNA was amplified in a 7500 Real-Time PCR system (Applied Biosystems) using IDT primers (**Supplementary Table 1**) and the TaqMan Gene Expression Master Mix (Applied Biosystems). RT-qPCR data were expressed as 2^−ΔΔCT^ relative to gene expression of cells in steady-state. ΔCT values were obtained using *Cdkn1a* as housekeeper gene.

### Western blot

Necrotic hepatocyte debris was generated from HepG2 cells by inducing mechanical disruption using a pellet mixer for 5 minutes in Hank’s Balanced Salt Solution (HBSS, Gibco, pH 8.5) supplemented with Ca^2+^/Mg^2+^ and 1X Halt Protease Inhibitor Cocktail (Thermo Fisher Scientific). The debris (from 1 x 10^6^ HepG2 cells) was treated with varying concentrations of peptides (PLK, PDK, PLE) ranging from 0.1 to 100 µM, Poly-L-Lysine (Sigma-Aldrich) ranging from 0.5 to 50 µg/mL, or heparin (LEO Pharma) ranging from 2.5 to 2500 U/mL, all prepared in HBSS. Incubations were performed for 2 hours at 37 °C. After treatment, the supernatants were collected and subjected to SDS-PAGE using 16% Tris-Glycine gels (Novex WedgeWell, Invitrogen, Thermo Fisher Scientific) with running buffer (25 mM Tris, 192 mM glycine and 0.1% SDS, pH 8.6). 10 µg of HepG2 full lysate, generated using RIPA Lysis and Extraction Buffer (Thermo Fisher Scientific), was loaded as positive control. Proteins were then transferred to a PVDF membrane and blocked with 5% BSA in washing buffer (20 mM Tris, 150 mM NaCl, 0.1% Tween 20) for 1 hour at room temperature (RT) under constant agitation. The membrane was subsequently washed three times for 5 minutes with washing buffer. Following the washes, the membrane was incubated overnight at 4 °C with agitation, using either rabbit anti-histone H3 antibody (1:2000, Abcam, Cambridge, UK) or mouse monoclonal anti-β-actin antibody (1:2000, Sigma-Aldrich). After three additional washes, the membrane was incubated with secondary antibody (either goat anti-rabbit IgG IRDye 800CW or goat anti-mouse IgG IRDye 680RD; 1:10,000; LI-COR Biosciences, Lincoln, NE, USA) for 1 hour in the dark at RT under agitation. The membrane was then washed as aforementioned and imaged using an Odyssey Fc Imaging System 7 (LI-COR Biosciences) at 800 and 700 nm. The images were analyzed with Image Studio Lite software (LI-COR Biosciences).

### Debris spot

Necrotic hepatocyte debris was generated as described above. The debris spot was generated by adding 6 µL of debris suspension, *i.e.* debris from 60000 HepG2 cells, into an 8-well chambered coverslip (Ibidi, Gräfelfing, DE) and drying for 2 hours in the laminar flow. The debris spot was then blocked with HBSS supplemented with 10% FBS (Biowest, Nuaillé, FR) and 1% FcR blocking reagent (human) (Miltenyi Biotec, Auburn, CA, USA) for 15 minutes at RT. For histone and total actin staining, the debris spot was incubated with rabbit anti-histone H3 antibody (5 µg/mL) or mouse monoclonal anti-β-actin antibody (20 µg/mL) for 1 hour at RT. After three washes with HBSS, the debris spot was stained with Hoechst 33342 (10 μg/mL, Invitrogen), Alexa Fluor™ 488 Phalloidin (66 nM) and 10 µg/mL secondary antibodies Rhodamine Red™-X (RRX) AffiniPure™ Donkey Anti-Mouse IgG (H+L) and Alexa Fluor® 647 AffiniPure™ Donkey Anti-Rabbit IgG (H+L) (all 10 µg/mL, Jackson ImmunoResearch, West Grove, PA, USA) for 1 hour at RT. After another washing step, the hepatocytes debris spot was imaged using a Zeiss Axiovert 200M fluorescence microscope (ZEISS, Oberkochen, DE).

### Confocal intravital microscopy

Mice were prepared as described previously [42] and anaesthetized with a subcutaneous injection of 80 mg/kg ketamine and 4 mg/kg xylazine. Fluorescent dyes were prepared by dissolving them in 100 µL of sterile PBS, followed by i.v. injection 10 minutes prior to surgery. To visualize extracellular DNA, mice were given an i.v. injection of SYTOX™ Green Nucleic Acid Stain (10 µM, Thermo Fischer Scientific). For F-actin staining, Alexa Fluor™ 555/647 Phalloidin was injected (2.2 μM, Thermo Fisher Scientific). Additionally, 50 µg of fluorescently labeled DNase 1 (Alexa Fluor™ 594, Invitrogen) or bovine serum albumin [BSA, tetramethylrhodamine (TAMRA) conjugate, Invitrogen] were injected to visualize labeling of necrotic areas by DNase 1. *In vivo* microscopy was performed using an Andor Dragonfly 200 confocal microscope (Oxford Instruments, Abingdon, UK). The images were visualized with Imaris v9.9.1 Cell Imaging Software (Bitplane, Zurich, CH). Extracellular DNA was removed *in vivo* by administering 40 mg/kg DNase 1 i.v. For experiments involving tPa administration, mice were injected i.v. with 5 mg/kg tPa. The mean fluorescent intensity (MFI) of DNA and F-actin in necrotic areas was measured over time following this treatment.

Kupffer cell depletion was achieved by i.v. injection of 200 µL clodronate liposomes (Liposoma, Amsterdam, NL) 48 hours before the experiment. To visualize clearance by the reticuloendothelial system, 4 µg/mouse of Brilliant Violet 421™ anti-mouse F4/80 Antibody (Biolegend, San Diego, CA, USA) and PE anti-mouse CD206 (MMR) Antibody (Biolegend) were injected. Necrotic hepatocyte debris, generated as previously described, was labeled for at least 30 minutes with an excess (5 µg per 10 x 10^6^ HepG2 cells) of DyLight™ 650 N-hydroxysuccinimide (NHS) Ester (Thermo Fisher Scientific) in 0.1 M sodium bicarbonate at pH 8.4. The unbound DyLight™ 650 NHS Ester was removed by centrifugation (5 min, 13 000 x g, RT), whereafter the debris was washed. Debris generated from 500,000 HepG2 cells was injected i.v. Using Imaris software, surfaces were overlaid onto the cells and necrotic debris through thresholding, after which the volume of overlap was calculated in µm^3^. The MFI of the signal in necrotic areas versus blood vessels was quantified using Fiji software. Quantifications in intravital microscopy were performed with n ≥ 3 mice, with different fields imaged in each mouse.

### Liver cryosectioning and immunostaining

To conduct liver immunostaining, the left liver lobe was harvested, embedded in frozen mounting medium (Tissue-Tek® O.C.T. Compound, Sakura, Torrance, CA, USA), and snap-frozen by immersion in liquid nitrogen. Cryosections of 10 µm thickness were generated using a cryostat (Microm Cryo-Star HM560, Thermo Fisher Scientific) and placed onto Polysine™ Adhesion Microscope Slides (Epredia, Portsmouth, NH, USA). The sections were fixed with 4% paraformaldehyde (PFA) in HBSS (pH 7.2) supplemented with Ca^2+^/Mg^2+^ and 0.1% BSA (Carl Roth, Karlsruhe, DE) for 1 hour at RT. Then, the sections were washed with HBSS and permeabilized with 0.1% Triton X-100 in HBSS for 1 hour at RT. Livers sections were washed again and blocked using 10% FBS and 1% FcR blocking reagent (mouse) (Miltenyi Biotec, Auburn, CA, USA) in HBSS during 1 hour at RT. To visualize the necrotic areas in liver cryosections, the sections were stained for fibrin(ogen) [37] using a polyclonal rabbit anti-human fibrin(ogen) antibody (28.5 mg/mL in HBSS, Agilent Dako, Glostrup, DK) overnight at 4 °C. To assess proliferation, Ki67 labeling was performed by labeling with a rabbit recombinant monoclonal Ki67 antibody [SP6] (0.15 μg/mL, Abcam) overnight at 4 °C. For histone and total actin staining, sections were incubated with rabbit anti-histone H3 antibody (5 µg/mL) or mouse monoclonal anti-β-actin antibody (20 µg/mL) overnight at 4 °C. The sections were washed and incubated with secondary antibodies for 2 hours at RT: Alexa Fluor® 488 AffiniPure™ Donkey Anti-Rabbit IgG (H+L), Rhodamine Red™-X (RRX) AffiniPure™ Donkey Anti-Mouse IgG (H+L) or Alexa Fluor® 647 AffiniPure™ Donkey Anti-Rabbit IgG (H+L) (all 10 µg/mL, Jackson ImmunoResearch). After another washing step, sections were incubated with Alexa Fluor™ 488/555 Phalloidin (66 nM) and Hoechst 33342 (10 μg/mL) for 1 hour at RT. A final wash was performed, after which mounting medium was applied (ProLong Diamond, Thermo Fisher Scientific). Images were acquired using an Andor Dragonfly Confocal Microscope or a Zeiss Axiovert 200M fluorescence microscope, and analyzed with Fiji. Stained areas were selected through thresholding, from which the percentage area of staining was determined.

### Flow cytometry of liver non-parenchymal cells

For the purification of liver non-parenchymal cells (NPCs), median liver lobes were harvested in RPMI 1640 medium (VWR, Radnor, PA, USA) and mechanically minced using the gentleMACS Dissociator (Miltenyi Biotec). To this suspension, 2.5 mg of collagenase D (Roche, Basel, Switzerland) was added for enzymatic digestion on a bench shaker for 1 hour at 37 °C. Subsequently, the liver homogenates were washed with PBS (300 x g, 5 minutes, 4 °C). The pellet was resuspended again in PBS, centrifuged at 60 x g for 3 minutes at 4 °C, and the supernatants containing the NPCs were harvested and filtered through a 70 µm cell strainer to remove undigested tissue. The cells were centrifuged again at 300 x g for 5 minutes at 4 °C, the supernatants were discarded, and the red blood cells were lysed with 1 mL of ACK Lysing Buffer (Gibco) for 10 minutes on ice. The ACK buffer was washed away by adding 4 mL of PBS and centrifuging at 300 x g for 5 minutes at 4 °C. The supernatants were discarded, and 1 x 10^6^ cells per condition were used for further analysis.

The cells were labeled with a viability dye marker (Zombie Aqua, Biolegend) and blocked with PBS supplemented with 1% mouse Fc receptor (FcR) blocking reagent (Miltenyi Biotec) for 15 minutes at 4 °C. The cells were then washed with 1 mL FACS buffer (PBS supplemented with 0.5% BSA and 2 mM EDTA), centrifuged at 300 x g for 5 minutes at 4 °C, and supernatants were discarded. Next, the cells were labeled with different antibodies [diluted in BD Horizon Brilliant Stain buffer (BD Biosciences, Franklin Lakes, NJ, USA)] (**Supplementary Table 2**) for 25 minutes at 4 °C in the dark. After labeling, the samples were washed by adding 1 mL FACS buffer, centrifuged at 300 x g for 5 min at 4 °C, and resuspended in 300 µL FACS buffer. Cells were immediately read in a Fortessa X-20 (BD Biosciences). The data were analyzed using FlowJo v10.8.1 (FlowJo, Ashland, OR, USA). The gating strategy is shown in **Supplementary Figure 6**.

### On-tissue microdigestion

Liver cryosections of 10 µm thickness were cut using a cryostat (Leica Biosystems, Machelen, BE) and mounted on Polysine™ Adhesion Microscope Slides. The slides were dried and warmed to RT under vacuum for 10 minutes. In APAP-treated samples, necrotic regions of interest (ROIs) were identified under 2X magnification and their coordinates were recorded. To digest the tissue, 20 µg/mL trypsin (Promega, Madison, WI, USA) suspended in 100 mM triethylammonium bicarbonate buffer was deposited onto the ROIs using a microdispensing system (i-TWO – 300p, M24You, Berlin, DE). The robotic system performed a sequence of operations, including washing the outer capillary orifice, aspirating trypsin, and depositing 30 droplets of 40 pL onto each ROI coordinate. This process was repeated for 20 passes, while ensuring uniform droplet size and maintaining a constant chamber humidity of 70% to prevent drying. Deposition was performed for 1 hour, resulting in about 144 nL trypsin deposited per ROI. After processing the first batch of 10 ROIs, the deposition step was repeated for another set of 10 ROIs. Following digestion, the slides were incubated at 37 °C in a sealed glass chamber humidified with a 1:1 methanol/water solution for 1 hour and then dried under vacuum for 15 minutes. To extract the digested peptides, the following solutions were used [43]: 0.1% TFA in water, 4:1 ACN/0.1% TFA in water, and 7:3 methanol/0.1% TFA in water. The tissue sections were quartered into 1 mm x 1 mm grids. For each grid, 14 µL of each solution was deposited over the digested spots, and the extracts were manually pipetted 10 times before recovering as much volume as possible at the last aspiration step. This extraction was repeated for each solvent system, yielding 28 µL of extract per spot. In cases where the spots are distant from each other and are not present in the same grid, the total volume was divided per spot and extraction was performed separately. The extracts were then collected, frozen at −80 °C and dried using a SpeedVac.

### Liquid chromatography tandem mass spectrometry (LC-MS/MS)

The dried extracts were reconstituted in 10 µL of 0.1% TFA in water and vortexed for 30 seconds. Desalting was performed using C18 ZipTips (Merck) following published protocols. Briefly, the tips were activated by repeatedly aspirating and dispensing 10 µL of ACN five times, followed by 10 µL of 0.1% TFA five times. The samples were loaded by aspirating 20 times, washed with 10 µL of 0.1% TFA by aspirating 10 times, and then eluted using 80% ACN/0.1% TFA by aspirating 20 times. After removing the organic solvent by lyophilization, the samples were reconstituted in 20 µL of 0.1% formic acid, and total peptide concentration was measured using a NanoDrop™ 2000 spectrophotometer (Thermo Fisher Scientific). All proteomic experiments were conducted using an Evosep One liquid chromatography system coupled to a trapped ion mobility spectrometry time-of-flight instrument (TIMS TOF Pro, Bruker Daltonik GmBH, Bremen, DE) via a nanoelectrospray ion source (Captive spray, Buker Daltonik GmBH). A total of 0.5 µg of each sample was loaded onto an EV1106 endurance column (15 cm × 150 µm, 1.9 µm) on the Evosep One. Peptides were eluted at a flow rate of 0.5 µL/min using the 30 samples per day (SPD) method available on the Evosep website. With this method, separation was performed using a 44-minute gradient. Mobile phases A and B were 0.1% formic acid in water and 0.1% formic acid in ACN, respectively.

### Proteomics data analysis

The raw files were exported and processed using MaxQuant v2.3.1.0. The files were searched using the target-decoy matching using the mouse UniProt database (accessed on 2022.05.30), with the false discovery rate set at 1% at both the peptide-to-spectra matches (PSMs) and protein levels. Trypsin specifically cleaving at the C-terminal to Arg and Lys was indicated and up to 2 miscleavages were allowed. Deamidation (Asn and Gln) and oxidation (Met) were set as variable modifications. Label-Free Quantification (LFQ) using the MaxLFQ algorithm, and Match Between Runs were used using default settings. The maximum precursor and fragment ion mass tolerances were set at default for a TIMS-DDA data type. Data analysis was performed using Perseus v1.6.7. The MaxQuant output tables were filtered to exclude reverse, contaminant and “only identified by site modification” assignments prior to further processing. LFQ intensities were log2-transformed and median-normalized row-wise, then filtered for at least 30% quantification events in at least one treatment group. Analysis of Variance (ANOVA) was performed and comparisons with p-value ≤ 0.05 were considered significant. The z-score of the proteins with significant comparisons were then obtained and used for hierarchical clustering using Pearson correlation as the distance metric.

### Statistical analysis

All other statistical analyses were performed in GraphPad Prism v10.1.2. Normality was assessed using the Shapiro-Wilkinson test. Significance between two groups was analyzed by Student’s t test or Mann–Whitney U test, and between multiple groups with one-way ANOVA or Kruskal-Wallis test. Differences were considered significant if p ≤ 0.05. Grubb’s test (extreme studentized deviate) was applied to determine whether extreme values were significant outliers from the rest. Data were represented as mean ± SEM.

## Conflict of interest statement

The authors have no conflicts to disclose.

## Funding

SS and SV hold PhD fellowships from the Research Foundation of Flanders (FWO-Vlaanderen; 1116922N and SB1S56521N, respectively). This work is supported by FWO-Vlaanderen Junior Research Grants (G058421N and G025923N), a KU Leuven C1 grant (14/23/143) and the Rega Foundation.

## Author contributions

SS and PEM conceptualized the experiments and prepared the manuscript. SS, JQ, CK, SV, ES, MLC, NP, NB and MSM performed the experiments and analyzed the data. JQ and GB provided technical support and contributed to experiment discussions. SS, MSM, PP and PEM interpreted and discussed the data. The manuscript was revised by all the authors.

## Data availability statement

The original contributions presented in the study are included in the article/Supplementary Material. The proteomics dataset generated and analyzed during the current study are available upon request. Further inquiries can be directed to the corresponding author.

## Ethics approval

The animal study was approved by the Ethical Committee for Animal Experiments from KU Leuven (registry number: P128/2021). The study was conducted in accordance with the local legislation and institutional requirements.

## SUPPLEMENTARY FIGURES

**Figure S1.**
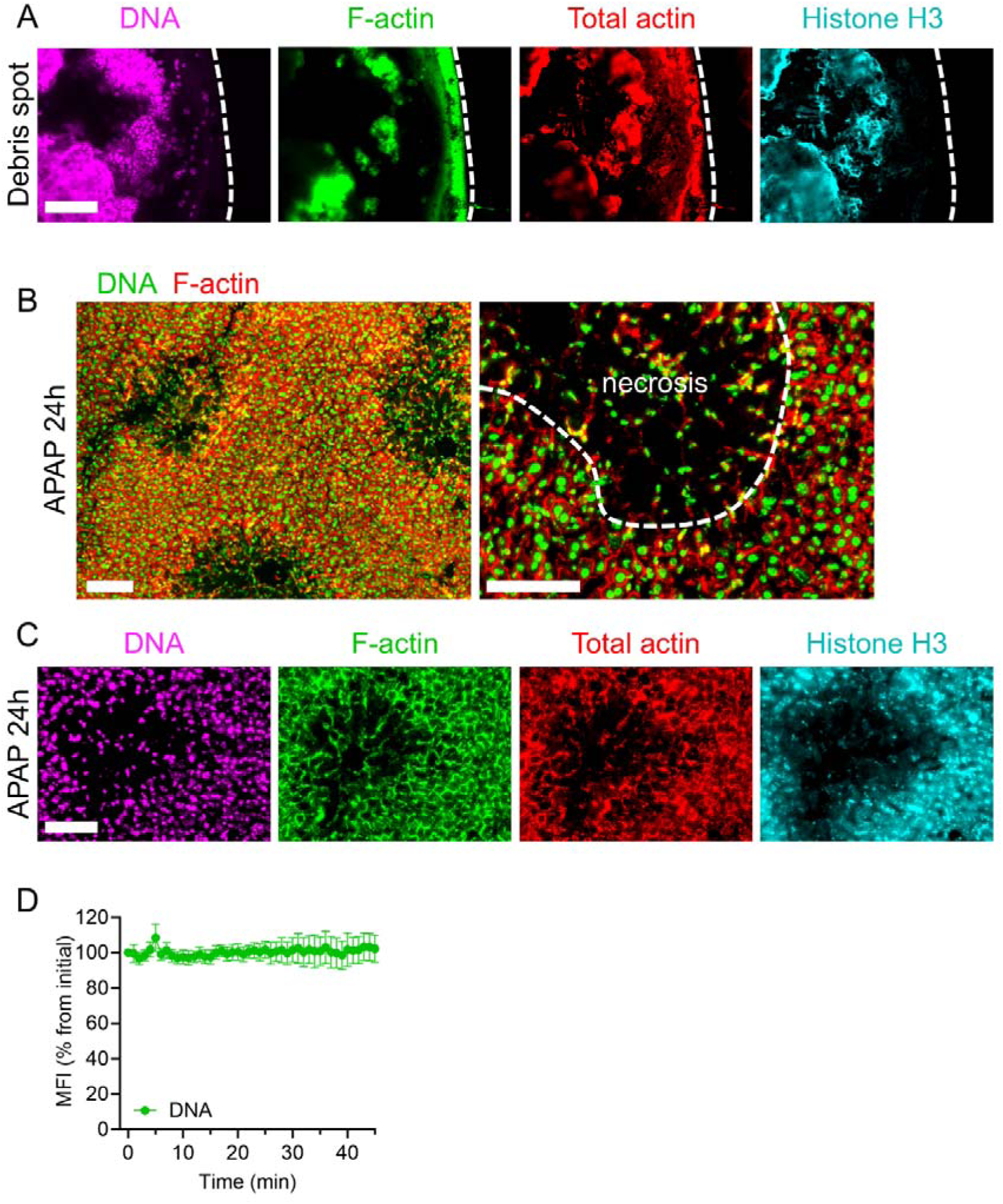
DNA, (F-)actin and histone labeling are absent in necrotic areas of mouse liver cryosections, but present in a necrotic hepatocyte debris spot. **(A)** *In vitro* hepatocyte debris spot stained for DNA (Hoechst, magenta), F-actin (Alexa Fluor 488 Phalloidin, green), total actin (anti-β-actin antibody, red) and histone H3 (anti-histone H3 antibody, cyan). Scale bar = 100 µm. Dotted lines indicate the edge of the debris spot. **(B)** Representative images of liver cryosections from mice 24 hours after APAP overdose (600 mg/kg) showing DNA (SYTOX Green, green) and F-actin (Alexa Fluor 555 Phalloidin, red) staining. Scale bars = 200 µm (left) and 100 µm (right). **(C)** Liver cryosections 24 hours after APAP overdose labeled as in (A). Scale bar = 100 µm. **(D)** Quantification of SYTOX Green mean fluorescence intensity (MFI) over time in necrotic areas during IVM, 24 hours after APAP overdose. Data are presented as mean ± SEM, pooled from 5 mice. APAP, acetaminophen.

**Figure S2.**
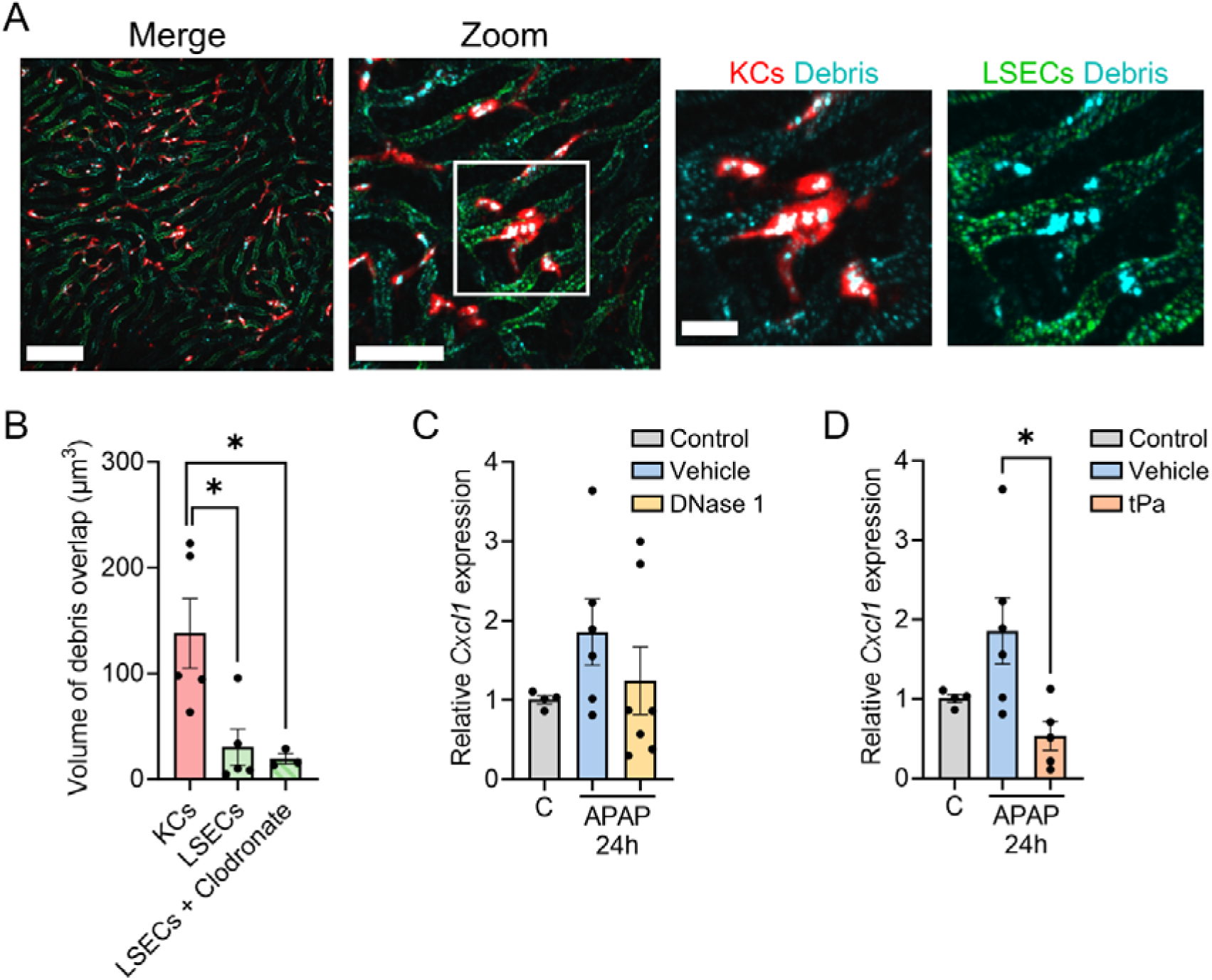
Debris capture by the reticuloendothelial system, and *Cxcl1* expression in livers of DNase- or tPa-treated mice. **(A)** Representative intravital microscopy images of healthy mice that were injected with fluorescently labeled (DyLight 650 NHS ester) necrotic debris generated from 500000 HepG2 cells. Kupffer cells (F4/80, red) and LSECs (CD206, green) are shown with debris overlap. Scale bars = 100 µm, 50 µm and 20 µm (from left to right). **(B)** Volume of debris overlap with Kupffer cells, liver sinusoidal endothelial cells (LSECs) and LSECs upon Kupffer cell depletion with clodronate liposomes. **(C)** Mice received an APAP overdose (600 mg/kg) and were treated i.v. with either vehicle (PBS) or DNase 1 (40 mg/kg) at 6 and 12 hours post-overdose for the 24-hour time point. *Cxcl1* expression levels in mouse livers, normalized to the average expression of a housekeeping gene (*Cdkn1a*), and presented as 2^−ΔΔCt^ relative to the control group. **(D)** *Cxcl1* expression levels in mouse livers of mice that received an APAP overdose and were treated at 12 hours with tPa (5 mg/kg) and were sacrificed at 24 hours. Each dot represents a single mouse. Data are represented as mean ± SEM. *p≤0.05 between indicated groups. C, control; APAP, acetaminophen.

**Figure S3.**
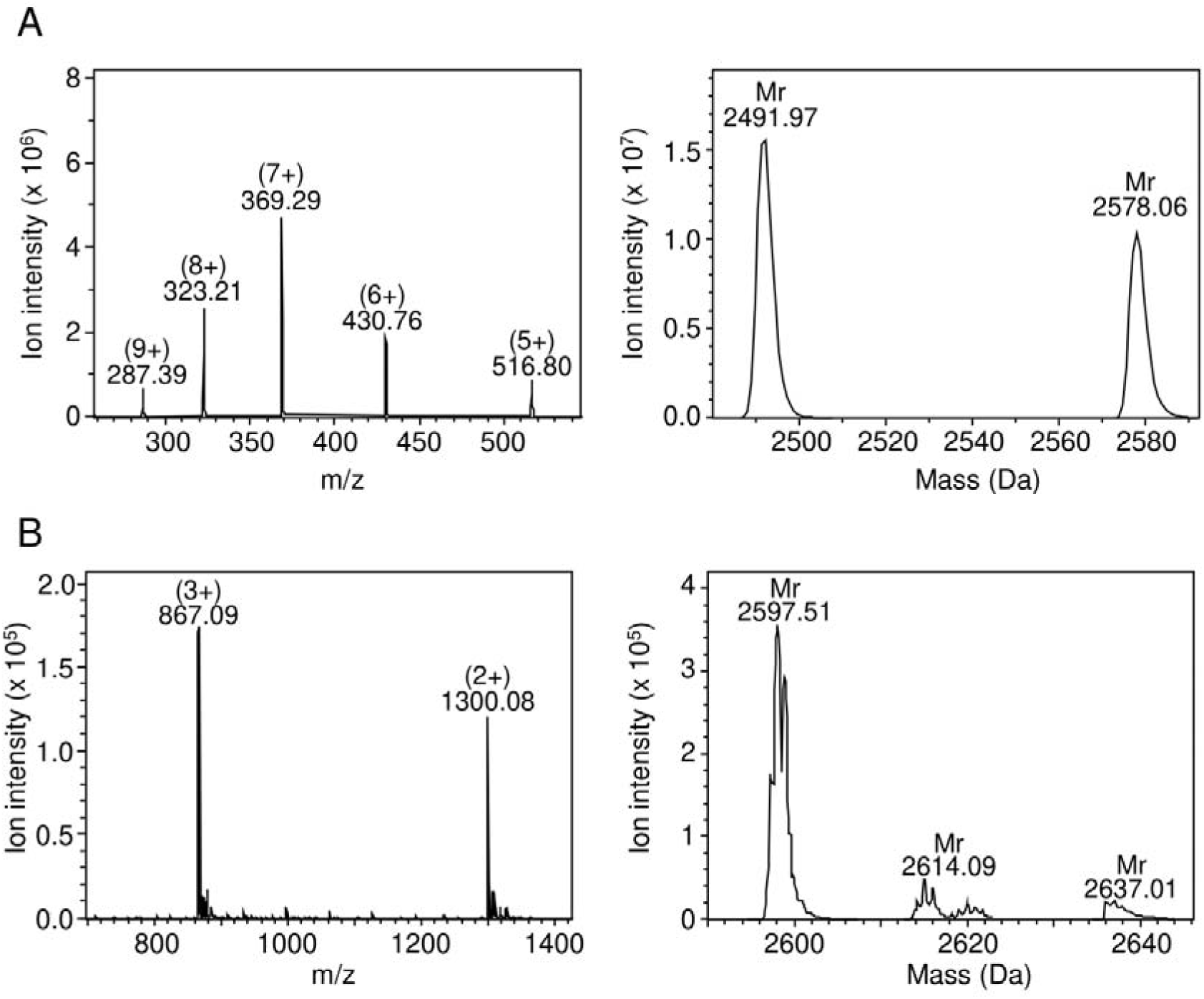
Identification of chemically synthesized peptides PLK and PLE. The peptides **(A)** PLK and **(B)** PLE were synthesized using Fmoc chemistry. After deprotection, RP-HPLC was used to purify the peptides and detection was performed using ion trap mass spectrometry. A mass spectrum of the peptide is shown with the intensity of the detected ions on the y-axis and the mass/charge (*m*/*z*) ratios on the x-axis. Deconvolution software (Bruker) was used to determine the relative molecular mass (M_r_) of the uncharged peptides. **(A)** The Theoretical Average Mr for PLK is 2580 Da and the Experimental Mr is 2578.06 Da. **(B)** The Theoretical Average Mr for PLE is 2599 Da and the Experimental Mr is 2597.51 Da.

**Figure S4.**
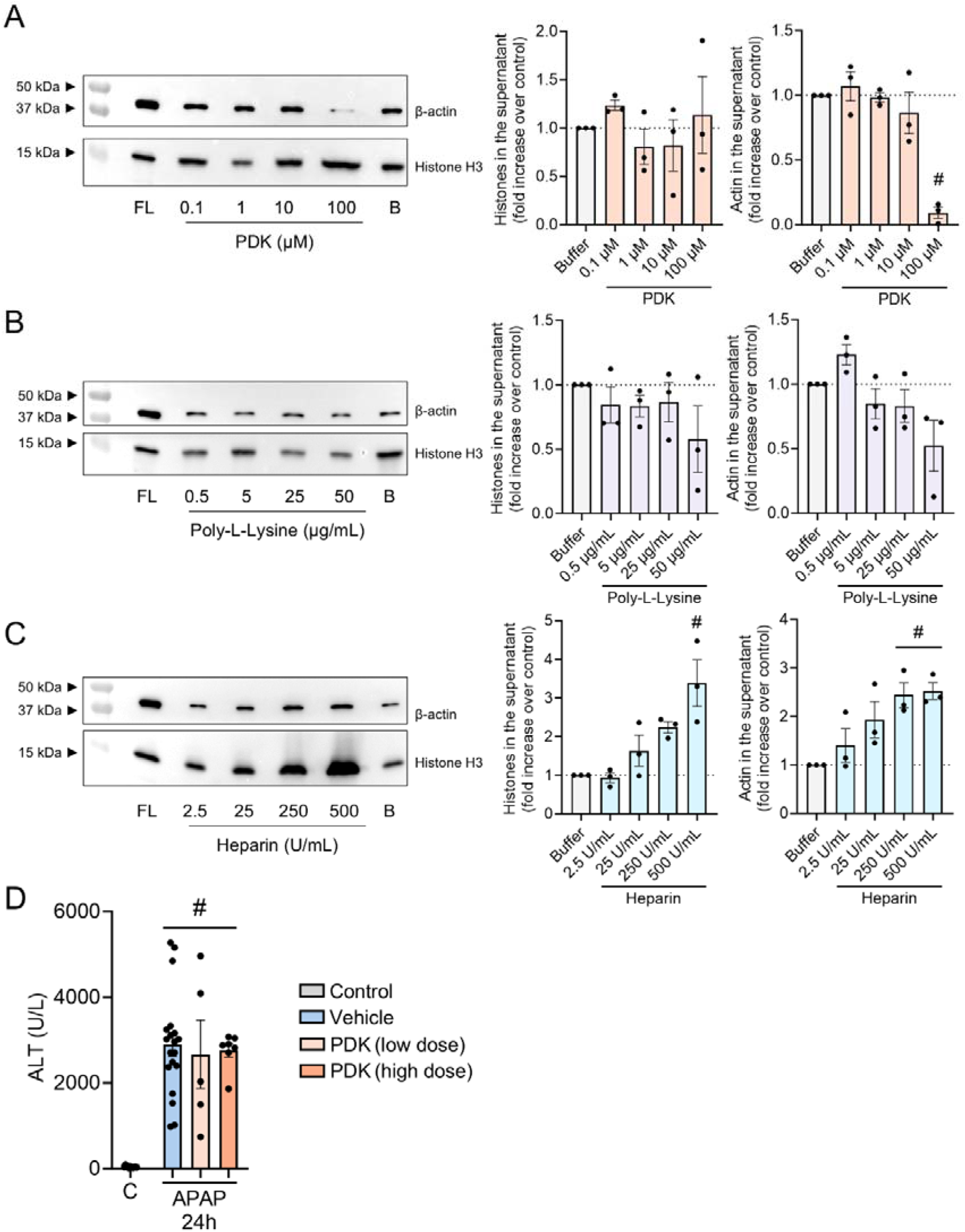
Efficacy of charged peptides and polymers in displacing necrotic debris *in vitro* and *in vivo*. Western blotting for histone H3 and β-actin in supernatants of HepG2 debris incubated with varying concentrations of **(A)** positively charged PDK, **(B)** positively charged Poly-L-Lysine, **(C)** negatively charged heparin, or buffer (B) alone. A full lysate of HepG2 cells (FL) was loaded at 10 µg as a positive control. **(D)** Serum alanine aminotransferase (ALT) levels in mice that received an APAP overdose (600 mg/kg) and were treated i.v. with either 400 µg/kg (low dose) or 800 µg/kg (high dose) of PDK at 6 and 12 hours, with analysis at the 24-hour time point. Each dot represents a single mouse. Data are represented as mean ± SEM. #p ≤ 0.05 compared to control. C, control; APAP, acetaminophen.

**Figure S5.**
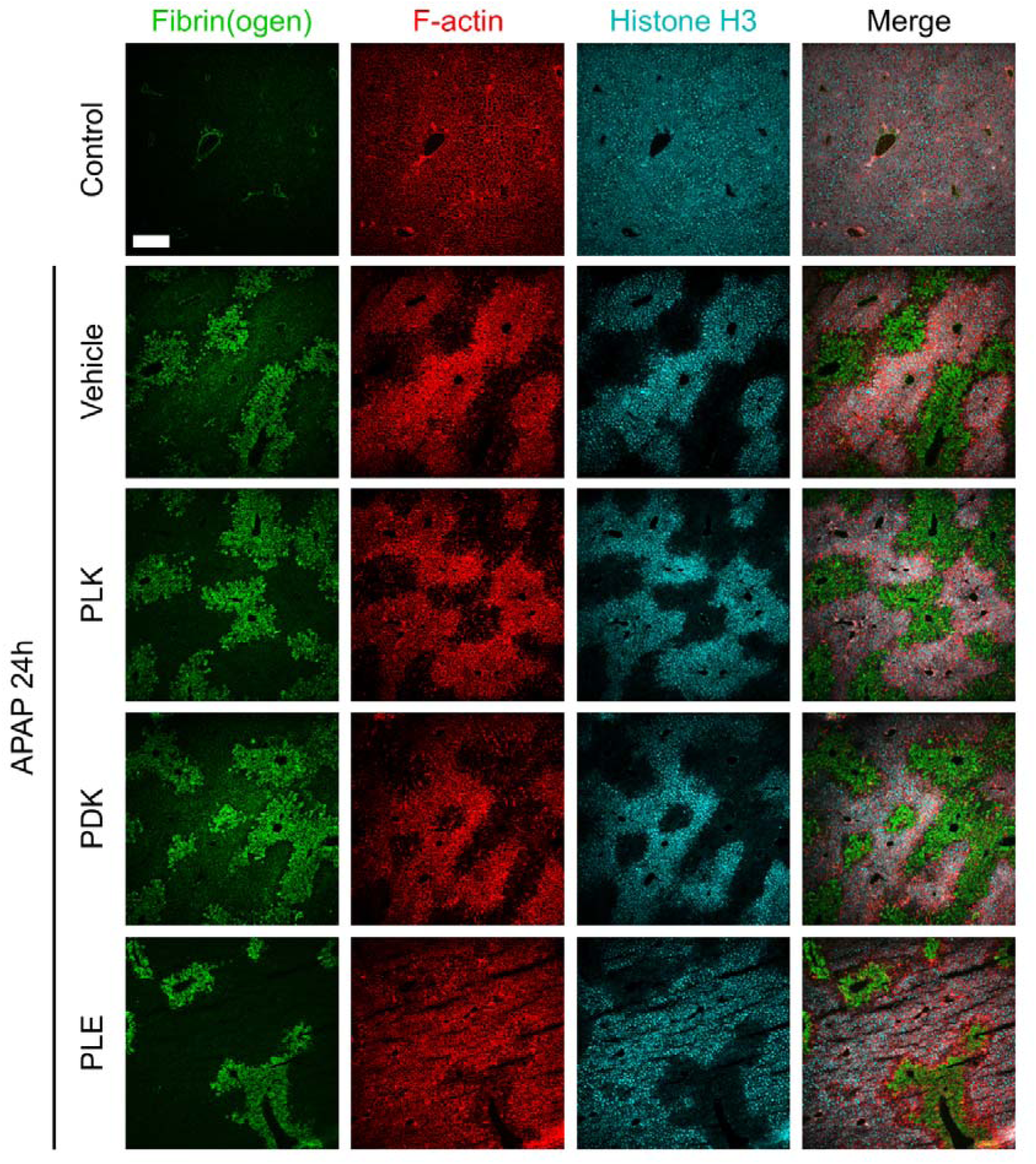
Fibrin(ogen), F-actin and Histone H3 labeling in liver cryosections of mice treated with peptides after APAP overdose. Representative images showing immunostaining of liver cryosections from APAP-challenged mice treated with vehicle (PBS) or peptides PLK, PDK or PLE (each at 800 µg/kg) and sacrificed 24 hours post-APAP. Green: fibrin(ogen); Red: F-actin; Cyan: histone H3. Scale bar = 200 µm.

**Figure S6.**
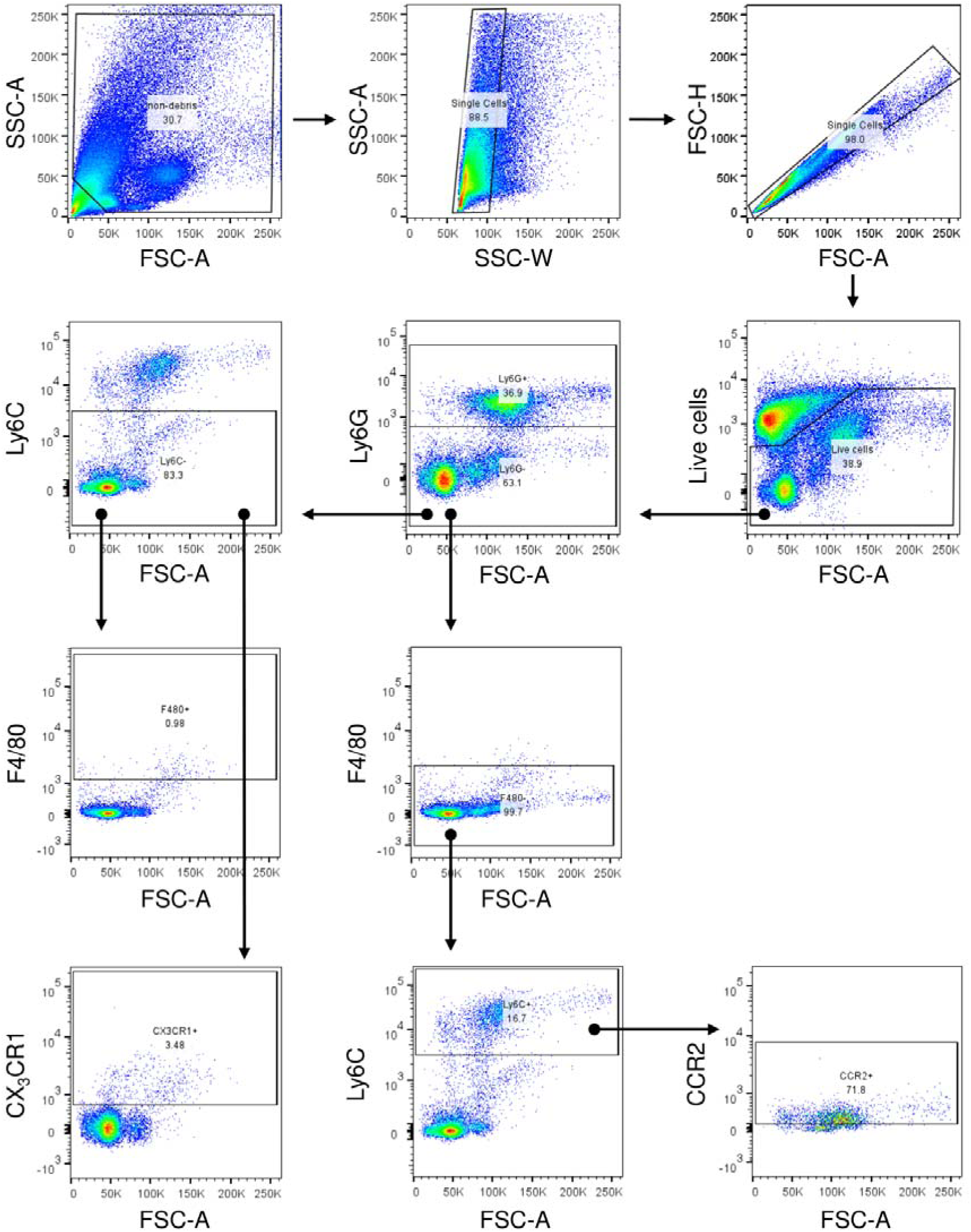
Gating strategy for liver non-parenchymal cells.

## SUPPLEMENTARY TABLES

**Supplementary Table 1.** Primers used for RT-qPCR.

| Name | Sequence | Supplier |
| --- | --- | --- |
| qPCR primer *Cdkn1a* | Mm.PT.58.5884610<br>Ref Seq: NM_007669 | Integrated DNA Technologies |
| qPCR primer *Il10* | Mm.PT.13531087<br>Ref Seq: NM_010548 | Integrated DNA Technologies |
| qPCR primer *Cxcl1* | Mm.PT.58.42076891<br>Ref Seq: NM_008176 | Integrated DNA Technologies |
| qPCR primer *Cxcl10* | Mm.PT.58.43575827<br>Ref Seq: NM_021274 | Integrated DNA Technologies |

**Supplementary Table 2.** Antibodies used for flow cytometry.

| Name | Supplier | Cat no. | Clone no. |
| --- | --- | --- | --- |
| Anti-mouse Ly-6G (Brilliant Ultra Violet™ 395) | BD Biosciences | 563978 | 1A8 |
| Anti-mouse F4/80 (Brilliant Violet 421™) | Biolegend | 123137 | BM8 |
| Anti-mouse Ly-6C (FITC) | Biolegend | 128006 | HK1.4 |
| Anti-mouse CX<sub>3</sub>CR1 (Alexa Fluor™ 647) | Biolegend | 149004 | SA011F11 |
| Anti-mouse CCR2 (Alexa Fluor™ 700) | R&D Systems | FAB5538N | 475301 |

## Notes

### Competing Interest Statement

The authors have declared no competing interest.

